# Causality organizes episodic memory and neural reinstatement

**DOI:** 10.64898/2026.07.22.740141

**Authors:** James W. Antony, Sara Abbas, Matthew S. Babb, Zachariah Reagh, Charan Ranganath

## Abstract

Endel Tulving proposed that episodic memory is organized by spatiotemporal relationships, and current theories center the role of time and space in organizing memory. However, these theories are supported by paradigms that eliminate other forms of relationships, like causality. Here, we used fMRI to reveal the organization of neural representations that support episodic memory with a paradigm that decoupled time and causal relationships. Participants were scanned while watching and recalling a TV show featuring five temporally interleaved storylines. Behaviorally, the order of recalled events was best predicted by the strongest cause-effect relationship relative to temporal, spatial, or semantic associations. Neural similarity across different events in regions of the default mode network (DMN) reflected causal structure, and DMN patterns at event boundaries specifically reactivated prior, causally related events, suggesting causality allows us to stitch together related events across temporal gaps. Additionally, causal network distance between successive events predicted the strength of DMN pattern changes across event boundaries, suggesting causality also predicts stronger representational switching between unrelated events. Together, these findings suggest that the DMN supports the construction of the complex causal networks that play a central role in organizing how we understand and remember events.

## Introduction

In 2012, Pixar creator Emma Coats shared a key principle that her company had developed to construct riveting, interconnected narratives ^1^. Each story was built on a structure that began with some initial setting of context followed by a chain of causally linked events:

“Once upon a time there was… Every day…

One day…

Because of that,…

Because of that,…

Until finally…”

Coats’ formula accords with the views of narrative theorists who propose that a narrative is not just a succession of events, but a succession of events with causal connections ^2,3^. Although narratives are typically associated with stories and films, several lines of evidence suggest that real-world episodic memories are also told and understood in the form of narratives ^4,5^. Despite compelling reasons to think about the role of causality in episodic memory, however, more evidence to suggest that causal relationships shape neural substrates of event comprehension and retrieval is needed ^6,7^.

Behavioral and neural research on free recall of word lists have reported considerable evidence to suggest that episodic memory may be organized according to temporal contiguity, or clustering the contents of recall by the time they were experienced ^8–12^. However, there is reason to think that temporal context cannot be the only factor for structuring episodic memory. Other factors like semantic relationships among stimuli ^13–15^, spatial context ^16^, reward states ^17,18^, and different instructions and rules ^15,19^, have also been shown to play important roles, sometimes eliminating the effects of temporal context altogether ^20^.

In narratives or other complex texts, several types of structure operate simultaneously – including time, place, characters, objects, and causality ^21^ – which naturally overlap. Critically, when researchers have disentangled time and causality by presenting scrambled or non-linear narratives, participants tend to mentally unscramble and recall events in a causal or chronological order ^22–26^. Moreover, when we directly contrasted evidence for several factors including time and causality, memory transitions were best captured by the strongest cause-effect (top causal) relationships ^25^. Therefore, causality appears to play an important role in memory organization for narratives ^27^ – possibly outpredicting the influence of time and other factors.

It is reasonable to think that causal inference might play a particularly important role during transitions between events, or “event boundaries.” According to theories of event cognition, when one event ends, we must construct a new mental model ^21,28–30^ of the current situation (“event model”) in order to understand the next event ^31,32^. Importantly, creating an accurate model of the current event often requires referencing (and thereby reactivating) prior events ^32,33,33–36^, effectively stitching them together. Therefore, one perceives current events in the context of a larger hierarchical thread of connected events ^27^, and we propose that these connected threads best reflect a series of causal inferences.

Neurally, leading candidates to reflect causal structure are regions of the default mode network (DMN). The DMN plays a critical role in integrating information across memories ^37,38^, represents abstract task states and narrative elements during encoding and recall ^29,39–41^, and reflects common inferences made across participants ^42–44^. Moreover, DMN regions specifically represent events in scrambled or temporally interleaved narratives ^7,45–49^, including high-comprehension, causally related “aha” moments, when the network reconfigures ^45^ and reflects patterns of prior related events ^7^. We will focus analyses on three parts of the DMN within the posterior medial system ^37^: the medial prefrontal cortex (mPFC), which plays a role in inferring latent causes ^50,51^, integrating old and new information ^52,53^, and representing hierarchical schemas ^39,41,54,55^ and other abstract structures and states ^56,57^; the angular gyrus (AG), which plays a prominent role in imagination and memory retrieval ^58–63^ and a causal role in linking information across events ^64^; and the posterior medial cortex (PMC), which plays a role in causal reasoning ^65^ and narrative comprehension and recall ^40,66–70^. Interestingly, during the continuous activation of an event model, neural activity has been shown to be relatively persistent ^39,71–73^, after which it rapidly switches to a new pattern ^39^. These “neural event boundaries” align with behavioral markers like agreement on where event boundaries exist across participants ^39,41,57,74–90^. Notably, event boundaries are not necessarily all-or-none but can be graded, which has been reflected behaviorally ^91^ and neurally ^57,74,92^. Therefore, DMN regions may conduct both the switching and stitching necessary for creating neural representations of complex causal networks during narrative comprehension.

Here, we investigated the impact of narrative causality on memory organization and neural substrates. We had participants undergo MRI scanning while watching and recalling a two-part episode of *Friends*, which had five (mostly) distinct, interleaved storylines (Figure 1). Another group of participants rated cause-effect relationships between each pair of events (Figure 1A), from which we determined the causal network structure (Figure 1C). Additionally, two groups of participants segmented the stimulus to validate analyses on neural event state transitions (Figure 1A). We found that the top cause-effect relationship for each event best predicted transitions during memory recall, demonstrating that causality has a profound effect on memory organization. Neurally, pattern similarity across events in the DMN reflected the causal network structure, and prior causally related events became reliably reactivated at event boundaries. Finally, neural event state transitions were stronger when there was a greater distance between successive events within the causal network. Therefore, causal network properties predicted memory recall and measures of both switching and stitching within DMN regions, suggesting causality plays a central role in narrative comprehension and memory.

**Figure 1.**
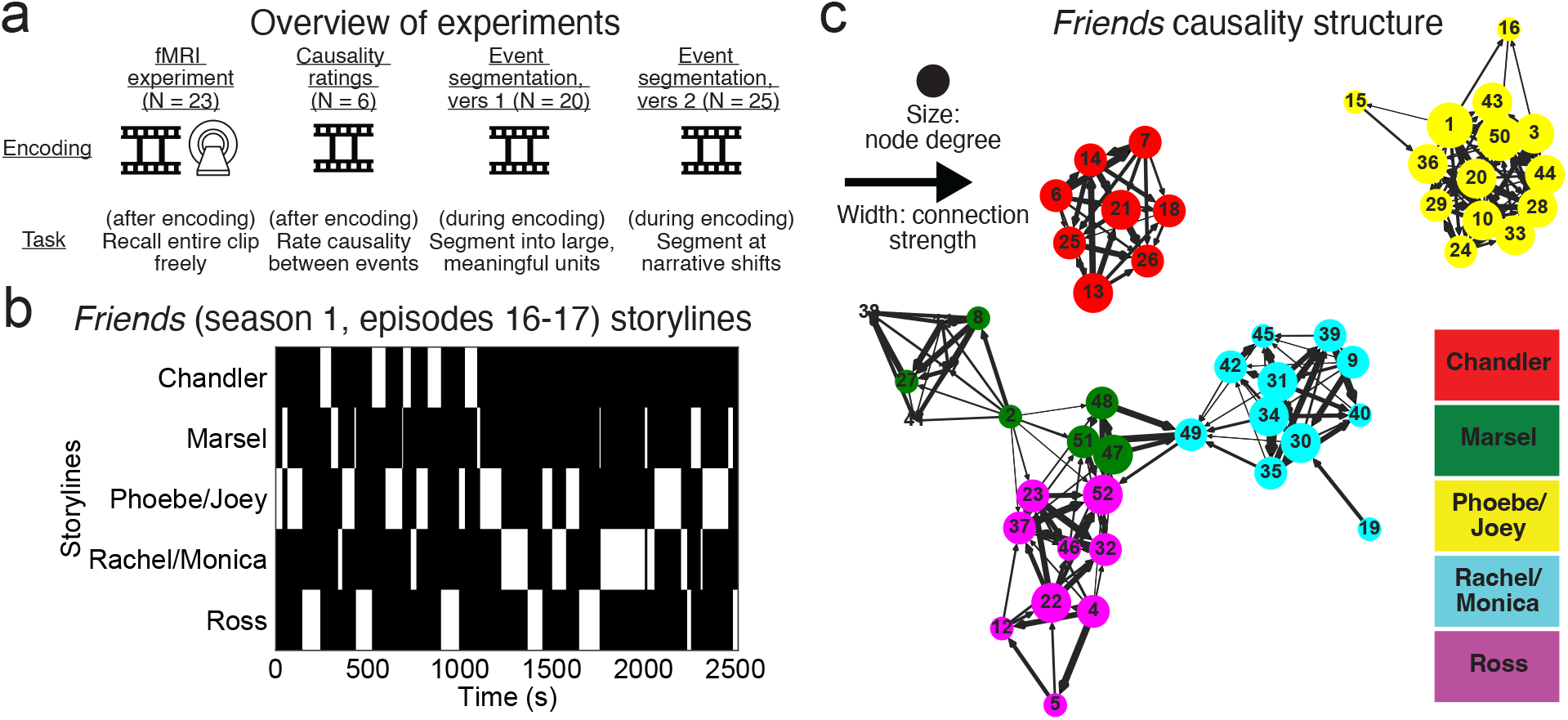
Overview of experiments, storylines, and causality structure. (a) Four groups of participants viewed a two-part episode of *Friends* under different conditions: (left) viewing during fMRI scanning, followed by free recall; viewing followed by rating the causality between each event pair (2nd from left); or viewing while marking segments between events according to meaningful units (2nd from right) or narrative shifts (right). (b) Time course of five interleaved storylines in the *Friends* stimulus. For time course of characters and places, see Figure S1. (c) Causality structure across events in graphical form ^27,93,94^.

## Results

### Causality between events

Participants (*N* = 6) who were not involved in the fMRI study were asked to view the *Friends* stimulus and rate from 0-10 how much each event causally affected each other event. We then averaged these ratings to create a causality matrix (Figure S2), from which we created a network plot of relationships between pairs of events (represented as edges between nodes) (Figure 1C). We also computed network measurements like causal network centrality and weighted path length between successive events (see Methods).

### Causal structure predicts memory organization and event memorability

Participants in the fMRI study (*N* = 23) were scanned during episode viewing and also during a test phase in which they were instructed to freely recall the episode in as much detail as possible. We first investigated factors that determine the memorability of a particular event, or the proportion of participants recalling each event, with particular interest for properties based on semantic networks (or those based on similar meaning across events) and causal networks. Prior work suggests that the density of semantic ^6,25,95^ and causal ^6,25,30,96–98^ relationships predict event memorability. Here, we defined semantic networks using the cosine similarity between events using the Universal Sentence Encoder ^99^) applied to the *Friends* screenplay (Figure S3). Supporting prior findings, we found that both causal and semantic network centrality, as measured by node degree for each event, or the weighted sum of edges to and from a given node, predicted memorability (causal: *r* = 0.38, *p* = 0.006; semantic: *r* = 0.42, *p* = 0.002) (Figure S4).

While both semantic and causal networks predicted what participants remembered, our primary interest was in how memories were organized, which we analyzed by measuring how participants transitioned while recalling successive events (Figure 2A). From these transitions, we computed and visualized the average likelihood of recalling a given next event based on the current event (Figure 2B). We next created matrices of several possible recall strategies to gain deeper insight into their relative influence (Figure 2C). These involved transitions between events based on time (presented order), characters, places, semantic relationships, or causal relationships. Additionally, we added strategies corresponding to the top semantic and causal relationships, whereby the strongest next event (given the current event) was assigned 1 and all others were assigned 0. Notably, while this binarizing step could remove useful information, it could also serve as a stronger predictor, given that each recalled event will transition to only one of the many remaining events that they could potentially sample from memory. Then, we examined the average evidence for each strategy at each transition. Critically, we found more evidence for the top causal strategy than all others (*p* < 0.001 for all comparisons; for correlations among all strategy models, see Figure S5). Therefore, while causality and semantics both determine what we remember, causality appears particularly important for determining how we organize memory.

**Figure 2.**
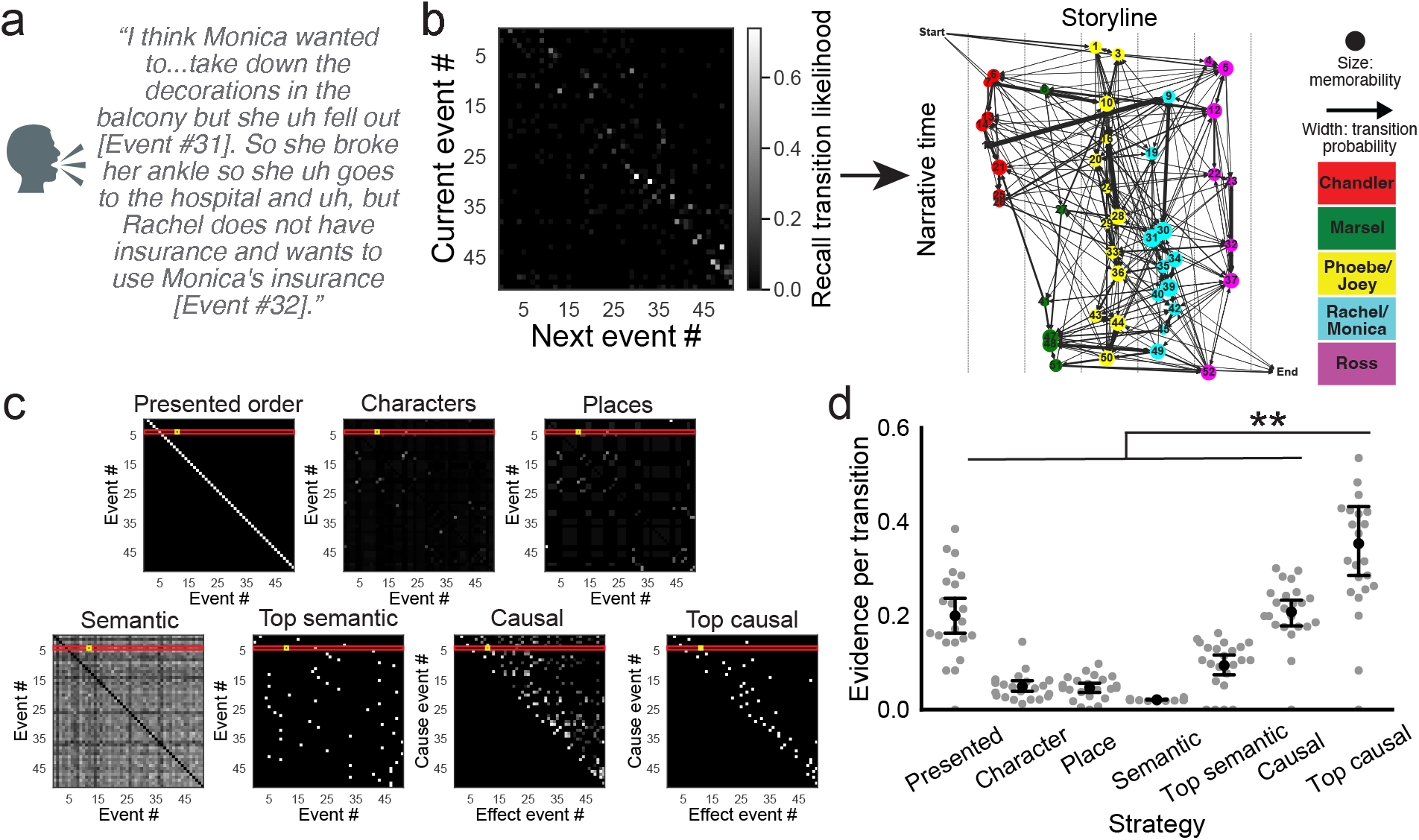
Causality predicts event memorability and memory organization during free recall. (a) Sample recall transition between two events. (b) Event recall transition probabilities between current and next recalled events (left). This transition matrix was used to create a recall network plot, with transition likelihoods shown via edge width between each labeled event on a plot organized by time within the narrative (*y*-axis) and storyline (columns within the *x*-axis) (right). This plot depicts how often participants transition within a storyline (within-column, near-vertical arrows), across storyline to nearby points in time (across-column, near-horizontal arrows), or some other strategy. (c) Transition matrices for various ideal recall strategies. (d) Mean evidence for each ideal strategy in (c) shows that the top causal model predicts memory organization better than all others (*p* < 0.001, all). For (d), gray dots indicate individual participants, black dots the mean, and black bars the standard error.

### Causal structure predicts neural event representational structure

Having established that causal relationships influence the memorability of events and the organization of episodic memory retrieval, we next investigated the degree to which causality, as opposed to other organizational factors, might account for neural representations of events. We hypothesized that the representational similarity of events would reflect not just their common storyline ^38,49^ but their graded causal structure, and we investigated this for each of six ROIs (Figure S6): V1 (to test the impact of visual features), three DMN regions part of the posterior medial system ^37^ [medial prefrontal cortex (mPFC), angular gyrus (AG), and posterior medial cortex (PMC)], and the posterior and anterior hippocampus for their disparate roles in memory ^100^ (postHC/antHC) [see Figure S7 for results in 100 cortical parcels ^101^]. We also modeled the influence of several other attributes that could influence similarity across events, including common characters ^46,55,102,103^, locations ^55^, and semantic features ^104^. Thus, any effect of causal relationships in this analysis would reflect unique variance over and above what could be accounted for by these other variables. Our approach involved first averaging TRs within an event for each voxel and correlating all event patterns with each other to create an event-by-event representational similarity matrix for each ROI (Figure 3A).

**Figure 3.**
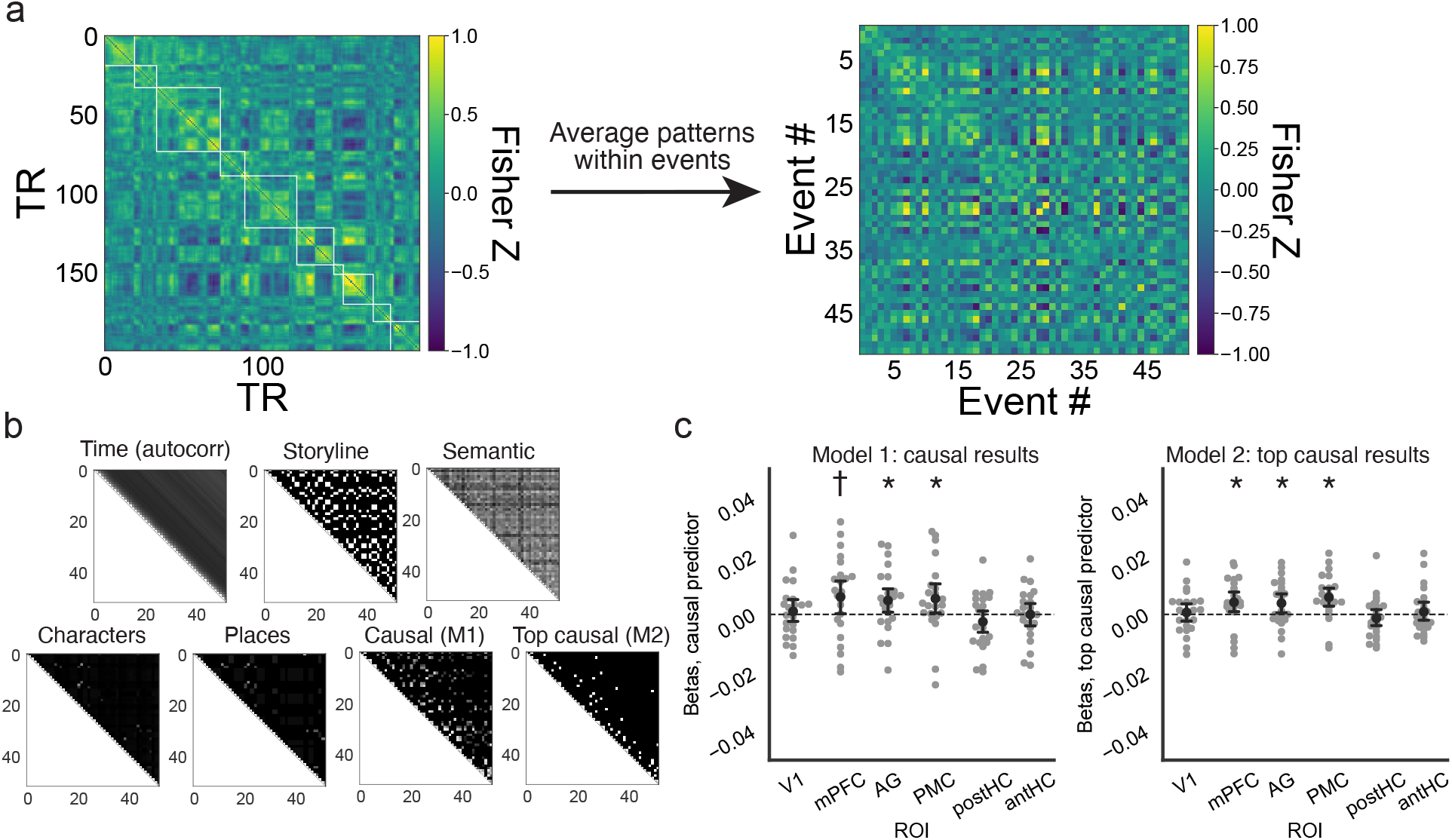
Neural event representations reflected causal structure. (a) (Left) Using data from the angular gyrus (AG), we plotted TR-by-TR representational similarity matrices for the first 200 TRs, with events outlined in white squares along the diagonal. (Right) Based on averaging patterns within events, we plotted event-by-event representational similarity matrices. (b) We used different predictors to model event-by-event similarity using multiple linear regression, from which we were primarily interested in the causal (Model 1 only) and top causal models (Model 2 only). (c) Betas for the causal predictor in Model 1 (left) and top causal predictor in Model 2 (right) showed that DMN regions reflect causal structure between events. For (c), gray dots indicate individual participants, black dots the mean, and black bars the standard error.

Next, we predicted the representational similarity for each ROI and participant using six regressors in two multiple linear regression models: time (based on the mean autocorrelated difference at each event lag), storyline [same storyline (1) from Figure 1B versus not (0)], semantic similarity, character similarity, place similarity, and either the causal matrix (Model 1 only) or top causal matrix (Model 2 only) (Figure 3B). Then we asked whether the participant-specific betas for the causal (Model 1) or top causal (Model 2) predictors statistically differed from 0.

We found that the three default mode regions had significant [AG: *t*(22) = 2.1, *p* = 0.044; PMC: *t*(22) = 2.1, *p* = 0.049] or marginally significant [mPFC: *t*(22) = 2.1, *p* = 0.050] relationships with causal structure (Model 1), and all three had significant relationships with top causal structure (Model 2) [mPFC: *t*(22) = 2.4, *p* = 0.02; AG: *t*(22) = 2.3, *p* = 0.03; PMC: *t*(22) = 3.6, *p* = 0.001]. V1, postHC, and antHC were not significant in either model [Model 1, V1: *t*(22) = 0.5, *p* = 0.60; postHC: *t*(22) = 1.2, *p* = 0.23; antHC: *t*(22) = 0.006, *p* = 0.995; Model 2, V1: *t*(22) = 0.4, *p* = 0.67; postHC: *t*(22) = 0.87, *p* = 0.40; antHC: *t*(22) = 0.57, *p* = 0.57]. Please see Figure S8 for results from all other predictors. Notably, prior studies have found storyline-related effects in the DMN ^38,48^, but we found unreliable effects of the storyline predictor in Models 1 and 2. However, another model (Model 3) with only time and storyline predictors showed that all three DMN regions predicted storyline [mPFC: *t*(22) = 2.9, *p* = 0.009; AG: *t*(22) = 4.3, *p* < 0.001; PMC: *t*(22) = 3.7, *p* = 0.001; Figure S9]. Collectively, these results suggest that causal structure predicts DMN patterns above-and-beyond the (binary) storyline, but that storyline predicts DMN patterns on its own. Therefore, neural representations were reflected in causal structure, suggesting that humans neurally stitch together causally related events during inference, offering a putative mechanism by which causality organizes recall.

### Prior, causally related events become reactivated at event boundaries

The preceding analyses showed how causality predicted the neural similarity between entire events, but recent findings show that prior events may be specifically reactivated at the boundaries between events ^35,36^. We next examined the extent to which patterns at event boundaries correlated with those from the non-boundary, middle parts of an event ^36^. To do this, we subtracted pattern matches between boundaries and past events (in which reactivation of prior events is possible) minus matches between boundaries and future events (Figure 4A-C). Here, we defined boundary patterns as the average activity of the boundary TR±2 TRs and the event middle pattern as the average pattern of 4 TRs after an event boundary to 4 TRs before the next event boundary (Figure 4A). We omitted events from ±2 events surrounding the current event boundary to avoid any issues with the autocorrelated BOLD signal bleeding into future events (Figure 4B). Overall, we found no significant differences between overall past minus future matches in any of our ROIs [V1: *t*(22) = 0.15, *p* = 0.88; mPFC: *t*(22) = 0.03, *p* = 0.98; AG: *t*(22) = 1.34, *p* = 0.19; PMC: *t*(22) = 0.69, *p* = 0.49; postHC: *t*(22) = 0.82, *p* = 0.42; antHC: *t*(22) = 1.31, *p* = 0.21] (Figure 4C). However, our prior behavioral and neural results suggested that reactivation might occur specifically for causally related events. Therefore, we also looked at the contrast between past minus future matches only for events with a non-zero causal relationship. Critically, we found significant differences in mPFC, AG, and PMC, but not in V1, postHC, or antHC [V1: *t*(22) = 0.14, *p* = 0.89; mPFC: *t*(22) = 2.50, *p* = 0.02; AG: *t*(22) = 3.58, *p* = 0.002; PMC: *t*(22) = 2.87, *p* = 0.008; postHC: *t*(22) = 0.70, *p* = 0.49; antHC: *t*(22) = 0.35, *p* = 0.73; Figure 4D]. Similar logic also led us to predict stronger matches for past events that were causally versus not causally related. In this analysis, we found significant differences only in PMC [V1: *t*(22) = 1.66, *p* = 0.11; mPFC: *t*(22) = 1.59, *p* = 0.13; AG: *t*(22) = 0.88, *p* = 0.39; PMC: *t*(22) = 2.72, *p* = 0.01; postHC: *t*(22) = 0.62, *p* = 0.54; postHC: *t*(22) = 0.86, *p* = 0.40].

**Figure 4.**
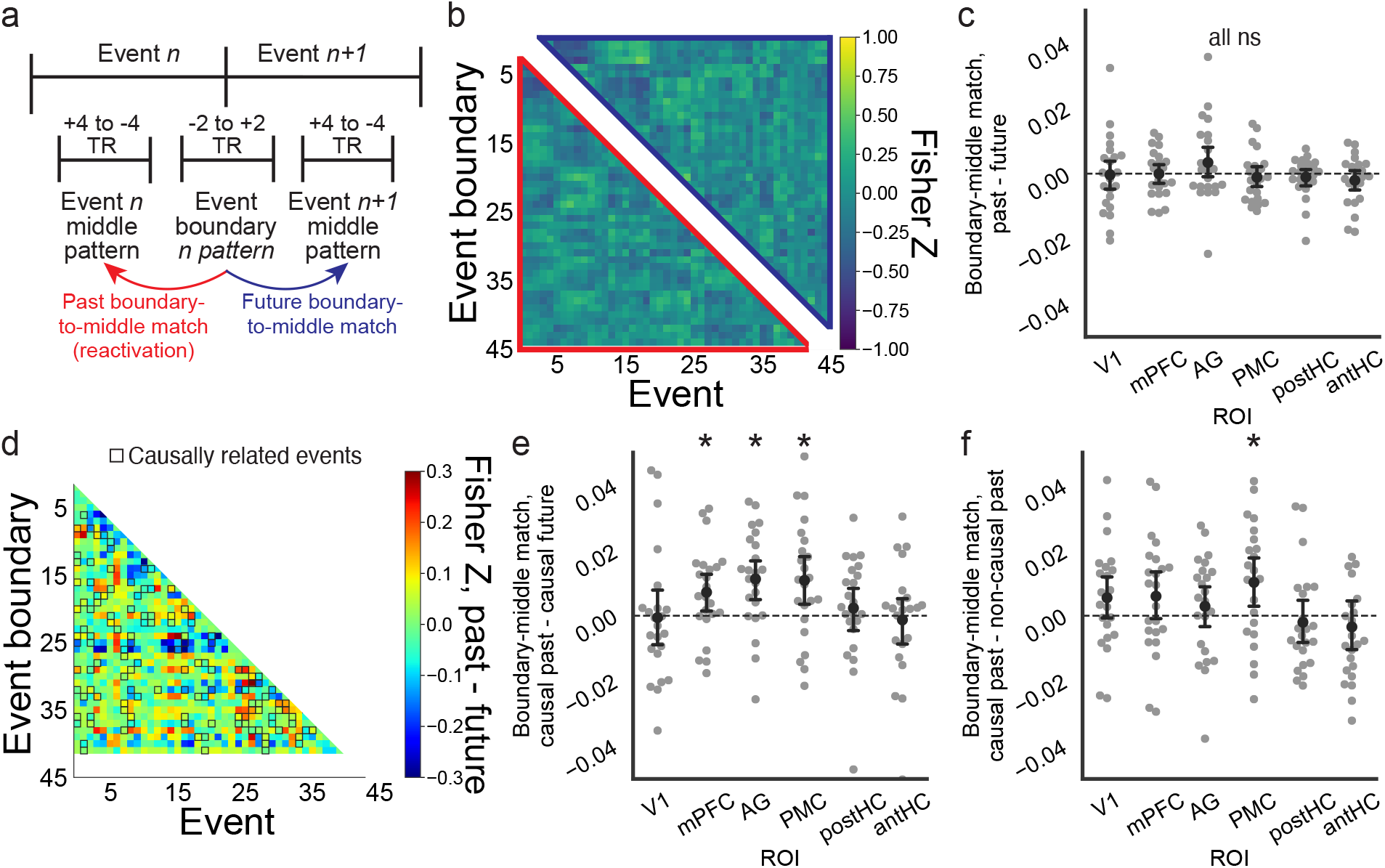
Causally related events become reactivated at event boundaries. (a) Analysis logic for contrasting between event boundary patterns with past versus future events ^36^. (b) Using data from the angular gyrus, the correlations (Fisher *Z* values) are shown between all event boundary patterns with patterns from the middle of past events (red) and future events (blue). (c) Contrasting boundary-middle matches for all past and future events produced no significant differences. (d) We plotted contrasts between past minute future events and highlighted causally related events (black outlines). (e) Contrasts between boundary-middle matches for past minus future events using only causally related events indicated significant differences in each DMN region. (f) Contrasts between boundary-middle matches for causally related minus not causally related past events similarly showed significant differences in PMC. For (c,e,f), gray dots indicate individual participants, black dots the mean, and black bars the standard error.

We next conducted additional and control analyses for these results. First, Hahamy et al. ^36^ used only the exact event boundary TR rather than the average of ±2 TRs, so we re-ran the analysis this way and similarly found mPFC, AG, and PMC had greater boundary-middle matches for causal past minus future events (all *p* < 0.03) and PMC had a greater boundary-middle match for causal minus non-causal past (*p* = 0.008). Second, we asked whether the number of TRs between each event boundary and event middle differed for any of the contrasts, and if so, whether this difference in temporal proximity could explain our results. There was no significant difference in comparing TRs between causal past versus causal future events [mean causal past TRs: 429.8±20.6, mean causal future TRs: 403.5±20.5, *t*(377) = 0.90, *p* = 0.37]. However, there was a difference between event boundaries and the middle of past causal and non-causal events, with causal event middle TRs being closer in time to event boundaries [mean non-causal past TRs: 495.2±9.2, *t*(1224) = 2.8, *p* = 0.005], suggesting that time could be a confounding factor in this analysis. To address this, we re-sampled the data 10,000 times by randomly selecting time points with replacement and examined the distribution of past causal minus non-causal differences only when there were no significant time differences (when the past causal vs. non-causal time difference contrast was *p >*= 0.5). In these cases, the distribution of boundary-middle match values for past causal minus non-causal events remained above zero 99.3% of the time in PMC (0.013±0.00005), suggesting the contrast remained when accounting for timing differences. Third, we conducted across-participant analyses by comparing the event pattern from one participant with the average event boundary pattern of all other participants ^36^. This analysis also produced significant differences in AG and PMC (but not mPFC) for causal past minus future pattern matches [mPFC: *t*(22) = 1.24, *p* = 0.23; AG: *t*(22) = 2.63, *p* = 0.015; PMC: *t*(22) = 4.82, *p* < 0.001] and PMC for causal minus non-causal past pattern matches [mPFC: *t*(22) = 1.30, *p* = 0.21; AG: *t*(22) = 0.26, *p* = 0.80; PMC: *t*(22) = 6.19, *p* < 0.001]. Finally, we also computed each of the above contrasts on 100 cortical parcels (Figure S10). Notably, there were no cortical parcels showing significantly stronger boundary-middle matches for all past minus future events. For the causal past minus future analysis, there was only 1 positive parcel (RH SomMot 8) after FDR correction, though there were 24 positive parcels (including 9 in the DMN) significant without FDR correction. For the causal minus non-causal past analysis, there were no parcels significant with FDR correction and 7 positive parcels (including 4 in the DMN) without FDR correction. In sum, our results (1) hold while using only the event boundary TR, (2) cannot be explained by timing differences between events and event boundaries, (3) hold in AG and PMC when using across-participant approaches (in which other auto-correlative signals should be minimal), and (4) generally hold, albeit less reliably, when using an across-cortical parcellation (rather than ROI-based) approach. Therefore, it appears prior causally related events become reactivated at event boundaries, suggesting event boundaries may provide a window even more temporally precise than full events in which prior and current experiences become stitched together.

### Alignment between behavioral and neural event boundaries

Event boundaries are commonly accompanied by rapid neural pattern changes ^39,57,74,105^, which have been proposed to underlie the understanding of a new event model ^29^. In this section, we will validate that there is internal consistency among our non-neural event boundary measures and show that they align with abrupt pattern shifts (“neural event boundaries”). There are multiple ways to operationalize event boundaries, including experimenter-defined boundaries or by computing boundary agreement in independent groups of participant raters. We used three measurements that could all theoretically align or predict neural event boundary changes, all of which we smoothed by ±1 s. The first was our experimenter-defined time course of event boundaries, which was a binary measurement based on whether there was an event change (1) or not (0) for each second of the stimulus. The second and third were average event boundary agreement values from two event segmentation experiments (Figure 1A). In the first experiment, participants demarcated events into subjectively large and meaningful units. We computed event boundary agreement (EBA1) from this time course, which correlated with experimenter-defined boundaries (*r*^2^ = 0.45, *p* < 0.001) (Figure 5A). In a second experiment (EBA2), we asked participants to demarcate events into moments when the narrative storyline changed. This EBA2 time course also correlated with experimenter-defined boundaries (*r*^2^ = 0.32, *p* < 0.001) (Figure 5A) and with EBA1 (*r*^2^ = 0.48, *p* < 0.001). Therefore, our event segmentation measurements aligned, suggesting each as a valid metric to investigate neural event boundaries.

**Figure 5.**
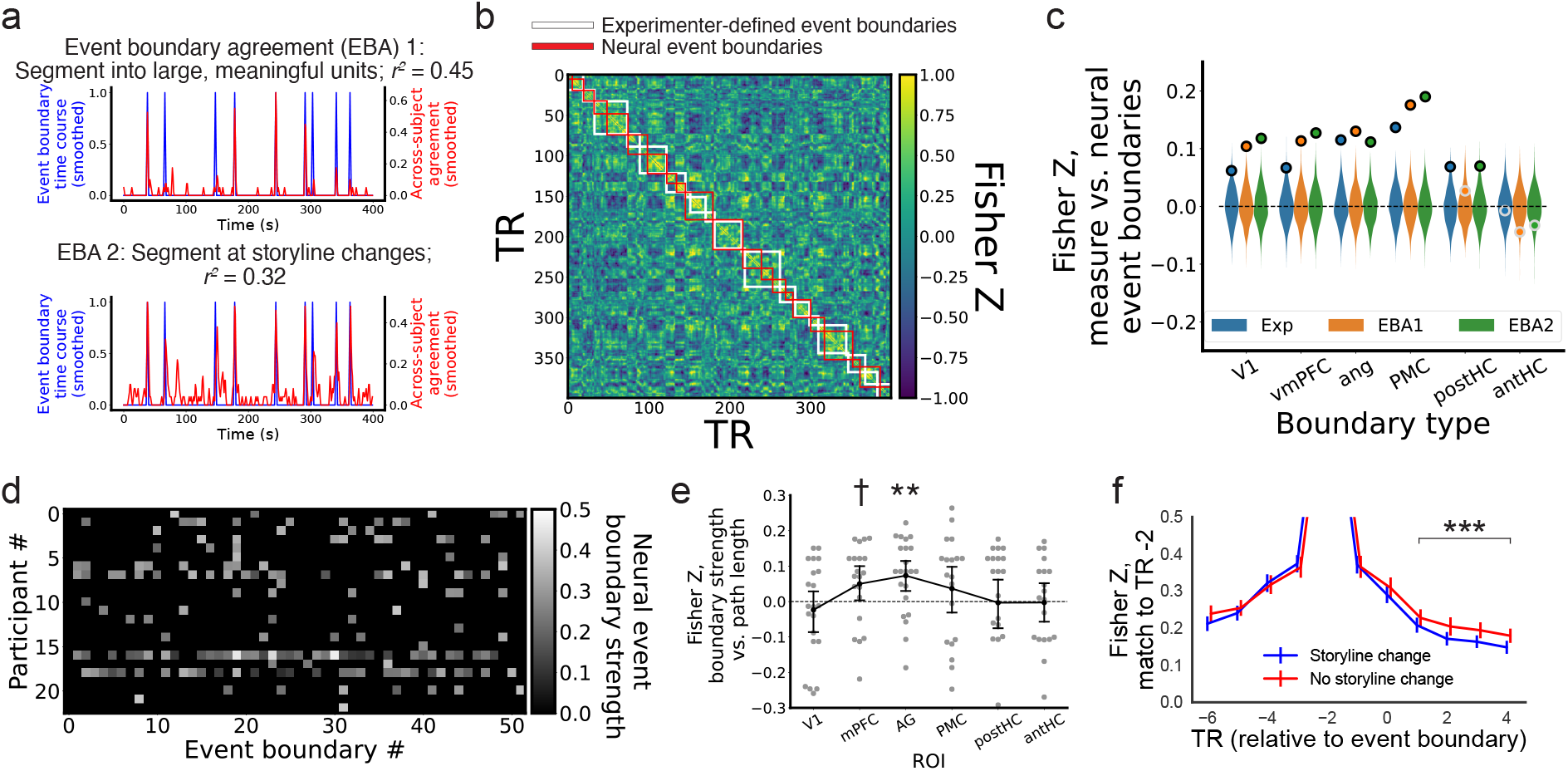
Neural event boundaries align with behavioral boundaries and correlate with causal network distance between adjacent events. (a) Alignment between experimenter-defined event boundaries and event boundary agreement in version 1 (EBA1; top) and version 2 (EBA2; bottom). (b) Sample TR-TR correlation matrix (across-participant average) for the angular gyrus against experimenter-defined boundaries (white) and neural event boundaries (red). (c) Alignment between neural event boundaries and experimenter- and group-defined boundary types across our primary ROIs. Circles with black boundaries were significant (*p* < 0.05), whereas those in gray were not. (d) Neural event boundary strength at each event boundary for every participant. (e) Correlation for each participant between event boundary strength for each ROI and weighted path length between successive events in the causal network. (f) Pattern change in angular gyrus before and after event boundaries depending on storyline change. For (e), gray dots indicate individual participants, black dots the mean, and black bars the standard error.

Next, we confirmed that these boundaries aligned with neural event boundaries derived via greedy state boundary search ^74^ (Figure 5B). Following prior approaches ^39^, we first reduced the neural dimensionality by applying an across-participant shared response model to the ROIs ^106^. We found the optimal number of features (dimensions) separately for each ROI by testing a range of features (5-40) and aligning neural event changes against experimenter-defined events from a video stimulus of another *Friends* episode (V1: 37; mPFC: 20; AG: 31; PMC: 29; postHC: 23; antHC: 5; see Methods). Next, using this optimal number of features applied to the main stimulus data, we first found time courses of each participant’s neural event boundary strength, or the correlation distance between successive neural state patterns, before averaging and smoothing these time courses across participants. Then we correlated this mean time course against each event boundary metric, and we assessed significance by contrasting the true value against the same correlations after scrambling the order of events. These alignments were significant in V1, mPFC, AG, and PMC (V1, exp-defined: *p* = 0.046; EBA1: *p* < 0.001; EBA2: *p* < 0.001; mPFC, exp-defined: *p* = 0.03; EBA1: *p* < 0.001; EBA2: *p* < 0.001; AG, exp-defined: *p* = 0.002; EBA1: *p* < 0.001; EBA2: *p* < 0.001; PMC, exp-defined: *p* < 0.001; EBA1: *p* < 0.001; EBA2: *p* < 0.001), sometimes significant in postHC (exp-defined: *p* = 0.02; EBA1: *p* = 0.29; EBA2: *p* = 0.018) and not significant in antHC (exp-defined: *p* = 0.78; EBA1: *p* = 0.10; EBA2: *p* = 0.23) (Figure 5C). Additionally, across-parcel results largely reflected the significant correlation between behavioral and neural event boundary types (Figure S11A-C). Therefore, we replicated prior findings ^39,57,74^ that behavioral and neural event boundaries largely align.

### Causal structure predicts neural event boundary strength

Rapid changes in neural event patterns (which underlie changes in neural event states) indicate reduced similarity between successive time points, providing a mechanism by which one could infer a new causal latent state ^34,107^. However, when one transitions to a new event in the same storyline (or one connected via an edge in the causal network), the neural pattern change should be weaker to reflect the similar underlying cause. Therefore, we reasoned that event changes should differ based on the distance between successive events within the causal network. To examine this, we found the weighted path length between successive events in the causal network, in which a strong direct connection would have a low path length value inverse to its causal edge strength (e.g., 0.11) and impossible connections were set to an arbitrary max value of 1. Then we correlated path length against neural event boundary strength for each participant in each ROI (see Figure 5D for an example from AG), and we asked whether the Fisher *z*-transformed correlations differed from zero. In this analysis, the AG was significant [*t*(22) = 3.80, *p* = 0.001], mPFC was marginally significant [*t*(22) = 1.92, *p* = 0.07], and V1, PMC, postHC, and antHC were not significant [V1: *t*(22) = 1.01, *p* = 0.32; PMC: *t*(22) = 0.96, *p* = 0.34; postHC: *t*(22) = 0.25, *p* = 0.81; antHC: *t*(22) = 0.55, *p* = 0.59] (Figure 5E). We also alternatively asked whether these results would change if, rather than event boundary strength, we used a binary time course based on whether any neural event boundary was present or not at experimenter-defined boundaries. We found similar results [V1: *t*(22) = 0.78, *p* = 0.44; mPFC: *t*(22) = 1.97, *p* = 0.06; AG: *t*(22) = 3.22, *p* = 0.004; PMC: *t*(22) = 1.06, *p* = 0.30; postHC: *t*(22) = 0.09, *p* = 0.93; antHC: *t*(22) = 0.10, *p* = 0.92]. Finally, an alternative way of testing this idea is to separate event boundaries depending on whether a storyline change occurred and then contrasting the correlation between a pre-event TR pattern (2 TRs before the event boundary) and post-event TRs (averaging TRs 1-4 after the boundary). This analysis produced a significant result in AG [*t*(22) = 3.5, *p* = 0.002; shown in Figure 5F] as well as mPFC [*t*(22) = 2.22, *p* = 0.037] and PMC [*t*(22) = 4.06, *p* < 0.001], but not in V1 [*t*(22) = 0.78, *p* = 0.44], postHC [*t*(22) = 0.68, *p* = 0.50], or antHC [*t*(22) = 1.42, *p* = 0.17]. Finally, across-parcel results showed one significant DMN parcel after FDR correction (LH Default Par 1) and four others significant without correction (RH Default PFCv 1, RH Default PFCdPFCm 3, RH Default pCunPCC 1, RH Default pCunPCC 2) (Figure S11D). Therefore, greater shifts between successive moments in the causal network resulted in greater DMN pattern shifts, suggesting a mechanism by which the brain can rapidly switch event models to separate adjacent, unrelated events.

## Discussion

Our findings demonstrate that causality plays a critical role in shaping memory and DMN states during comprehension of complex narratives. First, we found that the top cause-effect association was the best predictor for how participants transitioned during free memory recall of the narrative ^25^. Next, event patterns in the DMN reflected the narrative causal structure, and moreover, their event boundary patterns matched those from prior, causally related events, suggesting that event boundaries are an opportune time to reactivate and update prior memories. Finally, causal associations between successive events predicted the strength of pattern changes across event boundaries, suggesting that rapid neural changes reflected greater changes in mental inference. In sum, these findings suggest causality plays a central role in narrative comprehension and memory.

Memory transitions were best predicted not by time or semantics but by the event representing the strongest effect from a given cause. This finding may seem surprising for two reasons: (1) the prominence of time and semantics in prior investigations of memory organization ^8,15,108^ and inference processes ^109^ and (2) it involves eschewing potentially useful information (from other, weaker cause-effect relationships). Regarding (1), most prior investigations of recall organization have focused on lists of unrelated words, that by nature do not have causal links. In contrast, studies with narrative stimuli have shown that causality reliably predicts recall organization ^22–25^, suggesting that their omission from major memory models could be due to their absence in experiments removing more complex real-world variables. Regarding (2), prior theoretical work has suggested that free recall involves mentally reinstating not just a single past moment, but multiple moments simultaneously ^110^. Nevertheless, without any directive to do so, participants predominantly transitioned in recall using the top cause-effect relationship.

One interpretive framework for these results is that the causal network takes the form of a cognitive graph. Humans create complex cognitive graphs for spatial navigation, from which they can compute desired courses of action based on multi-step representations of successive states (successor representations) ^111–117^. In narratives, cognitive graphs could potentially be created along multiple dimensions in parallel (e.g., presented time, chronological time, place, characters, causality ^21^). Importantly, the development of a successor representation relies on transitioning from one state to the next during direct experience or memory reinstatement, the latter of which accords with our findings in that reactivating causally related events could link the two events. Therefore, just as humans use cognitive graphs or model-based transition structures to navigate through space or multi-step tasks ^112,115,117–119^, we propose that humans heavily rely on causal network representations to “navigate” through narratives.

Neurally, causal structure was reflected by DMN patterns across events. Prior work has shown that the DMN represents integration and inferences across memories ^29,37,38,43,49^, but there are many ways in which events can be linked. Here, we found that causal predictors significantly predicted DMN patterns after accounting for time, storyline, semantic structure, characters, and places. This similarity between causally related events suggests a mechanism by which individuals can reactivate and update prior event models depending on causality ^7,27,45,120,121^, which would lead to hierarchical threads of connected events important for understanding the full narrative. Intriguingly, a recent computational model trained to perform next-scene prediction preferentially retrieved causally related events while encoding new events ^122^, suggesting that linking causally related events aids narrative perception and prediction. In contrast to the DMN, postHC and antHC did not correlate with causality in Models 1-2, though the storyline predictor in postHC was significant in Models 1-2^49^. Prior findings, however, suggest that HC reinstatement occurs during retrieval in non-narrative paradigms ^123,124^ and that HC reactivation occurs at event boundaries ^36^ and helps bridge across experiences that fit into the same narrative ^35^. Therefore, future work is needed to determine the extent to which HC patterns reflect prior, reactivated events in narratives.

Analyses of the similarity between event and event boundary patterns suggested that prior events become reactivated and integrated in DMN regions like mPFC, AG, and especially PMC during current, causally related events. We did not find stronger event boundary-event middle matches for past than future events overall in hippocampus or in the main DMN ROIs, which contrasted with prior results following the same approach ^36^. However, those authors found stronger boundary-middle matches when the events were linked via semantic associations, which accords with our significant findings when specifically examining causally related events, in that meaningful associations between events predicted stronger reactivation at event boundaries. Together, our findings on across-event similarity and event-event boundary similarity in DMN regions suggest that prior, causally related events become reactivated and updated in the service of hierarchically linking events into causal chains to serve comprehension and recall.

Creating a causal network out of interleaved events also requires a mechanism to rapidly separate, or switch between, neural event patterns. A rich literature has uncovered both behavioral and neural effects demonstrating the consequences of such a mechanism. Behaviorally, event boundaries affect constructs like free recall or temporal order memory ^125–131^ and increased reading times ^132^ for information learned within versus across boundaries. Neurally, event offsets and boundaries result in univariate changes across the brain ^39,133–135^, and time courses of subjective or experimenter-defined event boundaries reliably correlate with multivariate pattern changes across the brain ^39,41,57,74,81,86,88,89,136^. Our neural event state results in DMN regions replicate these prior findings. Moreover, event cognition accounts suggest that an event boundary prompts switching to a new event model ^21,30^, which is either new or related to a prior event. Theoretically, when this occurs, there should be a larger shift when the boundary spans two events that are less causally related, which we corroborated by showing that neural event boundaries are significantly stronger in AG (and marginally significantly in mPFC) when there was a larger path length in the causal network between successive items. Therefore, DMN regions like mPFC and AG can conduct both the switching and stitching required to mentally construct a cognitive map of a causal network.

This study opens the door to future directions related to what types of content become reactivated and when. We did not examine event boundary-middle matches for the same event due to issues with the auto-correlated BOLD signal. Other studies have suggested that elements of the current event become reactivated at event boundaries ^83,137^, and moreover, recent computational modeling and fMRI findings suggest that selectively encoding information from the current event at the boundary – likely reactivating relevant information from the event – may be an optimal memory strategy from a resource-rational perspective ^138,139^ (though see ^140^). Additionally, other studies asked participants to press a button when they made an inference or later reported a subset of key moments from each event ^7,141^, both of which allow for zooming in on precise moments ripe for reactivation. We anticipate that studies using these tasks, perhaps combined with neural or intracranial EEG methods on event-based tasks ^142–146^, could produce highly time-resolved patterns to address when and how the stitching process underlying reactivation unfolds.

In summary, causality is critical for narratives – this is true for how artists create them ^1^, how narrative theorists define them ^2^, and, as we demonstrated, how narratives are remembered and represented in the brain. Our findings suggest that causality should be centered in conceptualizations of memory organization alongside (or even superseding) other factors like time and semantics. Moreover, they suggest that the DMN helps conduct the mental switching and stitching necessary for creating the complex meshwork underlying real-world memories.

## Methods

### Participants

In the fMRI experiment, participants (*N* = 23, 14 female, 19-29 years old) were recruited from the University of California, Davis community and screened for contraindications. Additionally, data from three other participants were lost due to technical scanning issues. In the causality ratings experiment, participants (*N* = 6, 4 female, 20-32 years old) were recruited from California Polytechnic State University, San Luis Obispo (Cal Poly) community and completed the study for course credit. In the event segmentation experiments, participants (Version 1: *N* = 20, 18-24 years old; Version 2: *N* = 25, 18-25 years old) were recruited from the University of California, Davis and completed the study for course credit. Data from 10 (Version 1: 8; Version 2: 2) participants were removed for confusion or non-compliance with instructions. All behavioral and fMRI participants at UC Davis gave consent according to UC Davis IRB #1352490. All Cal Poly participants gave their consent according to Cal Poly IRB #2020-068.

### Stimuli

Our main video stimulus was a two-part episode of *Friends* (1994) (season 1, episodes 16-17). The stimulus was concatenated and edited to remove time points that were irrelevant for our purposes, including the theme song, parts of the opening and closing credits, and long transitions between events (final version: 41 minutes, 59 seconds long). This stimulus was selected because it had (1) multiple (five) storylines with low causal influence on each other, and (2) multiple (23) instances in which storylines changed within a spatiotemporal scene (i.e., the conversation topic changed) and other (8) instances in which the same storyline persisted across spatiotemporal scene changes. Analyses related to the second reason are not featured in this paper.

To train shared response models (SRM) using an independent stimulus ^105,106^, in the final fMRI phase we showed participants a second clip from *Friends* (first 11:26 from season 1, episode 18). This video segment was only used for training the SRMs and was not recalled by the fMRI participants nor rated/segmented in other experiments, though we used experimenter-defined boundaries to examine the best neural event boundary alignment across a range of possible SRM dimensions.

### Causality and importance ratings experiment

Participants in this experiment watched the main stimulus, after which they received instructions for the causality and importance rating tasks. For the causality task, we gave them the following instructions ^25,147^: “Your job is to identify and make a list of event pairs that are causally related to each other within the TV show. How can we decide whether two events are causally related or not? In an extremely broad sense, one might say that any event that happened before a target event could be at least partially responsible for the event to happen (e.g., you were born because there was Big Bang), but this wouldn’t give us very useful information. So, we want to identify only those event pairs that are more strongly related, and you will need to use your own best judgment to decide whether the causal relationship is strong enough. For example, if we have a movie like below … Event 1: Jane orders a crab cake at a restaurant. Event 2: Jane finds a dead fly in her crab cake. Event 3: Jane complains to the manager of the restaurant. You may say that there is a stronger causal relationship between Event 2 and Event 3 than between Event 1 and Event 2. We don’t really have strict rules or criteria, so it is up to your subjective judgment. But please try to keep your criteria as consistent as possible.” We did not specify further dimensions along which causality should be inferred ^93,148^. Along with these instructions, they received a 2-D spreadsheet with a brief description of each event on each row and column, and they were asked to rate the causal influence of the events in each row on events from that column (from 1-10).

We quantified the reliability of the causal ratings in two ways. The first was by averaging all raters except one and correlating this N-1 average pattern with the remaining rater (range: Pearson *r* = 0.70 to 0.88, median *r* = 0.8). The second was to average pattern from the first three raters with the pattern from the next three raters (split-half reliability) (*r* = 0.88). We then used the mean ratings between each event pair as our causality matrix, with the exception that idiosyncratic cause-effect relationships (given a non-zero value by only a single rater) were assigned a 0.

We anticipate that future research will use automated, LLM-based methods to assess causality, which have thus far produced mixed but potentially promising results ^94,149–151^.

For the importance task, we gave participants the same event descriptions but this time asked them to simply rate the importance of those events in the context of the full episode (from 1-10). Note that importance correlated strongly with memorability across events (see below) (*r* = 0.73, *p* < 0.001), but we otherwise did not analyze it in this study.

### Semantic network and centrality measurement

To calculate semantic centrality, we first used the cosine similarity between embeddings from the Universal Sentence Encoder ^99^ applied to the *Friends* episode transcript. This produced a 2-D similarity matrix from which we created an undirected semantic network. Finally, we calculated the node degree (sum of edges connected to each node) as our measurement of semantic centrality ^147^.

### Event segmentation experiments

To obtain a behavioral measure of the segments between events, we collected data from two groups of participants who watched the same video stimulus. In the first version, they pressed a button according to the following instructions ^152^: “Your task today is to press the spacebar when, in your judgment, one unit of the show ends and another begins. Please mark off the show you’ll be seeing into the largest units that seem natural and meaningful to you.”

Following previous approaches with different event segmentation instructions ^134,153^, we also ran a second version of this experiment with different instructions: “Your task today is to press the spacebar when, in your judgment, there is a shift in the narrative or topic of conversation. Please mark off these shifts into the largest units that seem natural and meaningful to you.” Seconds of the movie clip were discretized according to the presence of a button press, with any press indicating a 1 and no press a 0. We also smoothed the mean time course by averaging the number of button presses from ±1 surrounding each second. Both sample sizes were larger than the *N* = 18 recommended previously to acquire stable representations of event boundary agreement ^154^. The experimenter-defined time course was similarly discretized by event boundary time courses at the level of the second, such that each second would either have a 1 (event boundary indicated) or 0 (no boundary). Next, we ran correlations between the smoothed experimenter-defined time courses and smoothed time courses of event boundary agreement (see *Alignment between behavioral and neural event boundaries and effects of causality* below).

### Free recall scoring and transition analyses

Research assistants transcribed and scored each 1-s segment of the audio recordings from the fMRI session, and they identified recall segments for specific events where applicable. To visualize recall transitions, we calculated the likelihood of recalling each next event from the current event by finding all instances of that event and all subsequent next events and summing that distribution to 1 within each current event (row) (Figure 2A). Next, to visualize transitions according to both time and storyline, we created a network plot with events as nodes and transitions as edges, adding starting and ending states. For this, we manipulated the *x*- and *y*-coordinates, such that the *x*-coordinate was consistent (with a minor jitter) within a storyline (in an arbitrary column order) and the *y*-coordinate charted presented order for each event (Figure 2B).

To investigate recall organization, we first created ideal transition matrices for several possible recall strategies ^25^. For all strategies, we considered the current event being recalled as the row, and the evidence within each column a prediction for the next event, so each strategy involved a probability distribution that summed to one within a given row. For the presented order strategy, the very next event was predicted in all cases. For the character strategy, because there were different numbers and combinations of characters present during each event, we found the characters involved in the current event and the combination with which they matched characters in each other event. For the place strategy, we included predictions to each other event occurring in that scene. For the semantic strategy, we used semantic similarity as described above. For the top semantic strategy, we binarized the semantic strategy matrix by making the highest similarity event within each row 1 and all others 0. For the causal strategy, we used average causality ratings (described above). For the top causal strategy, we binarized the causal strategy matrix by making the highest similarity event within each row 1 and all others 0.

To measure evidence for each strategy, we looped over recalled events in order for each participant. At each transition, we focused on the row of the currently recalled event and summed the strength of the evidence for the column of the next recalled event (Figure 2C-D). Last, we contrasted the strength of each strategy as within-participant *t*-tests between the top causal versus all other strategies.

### Network analyses

We created weighted network graphs using the Python ‘networkx’ toolbox ^155^. These graphs were directed for the causality network (from cause *→* effect) and recall network (from the current *→* next recalled event). The semantic network was undirected because one cannot infer directionality from Universal Sentence Encoder outputs. We calculated centrality as node degree, or the sum of all edge weights for a given node. We calculated weighted path length as the shortest path length between each pair of events, with pairs with no possible path set to 1^156^. Note that for visualization purposes, we only plotted the top 20% of edge weights, but all weights were used to calculate network properties.

To investigate network effects on memorability, we ran Pearson correlations on event memorability, or the proportion of participants who recalled each event, against causal and semantic network node degree values (Figure S4).

### fMRI experiment procedure

The fMRI experiment consisted of five scanned phases: (1) pre-movie picture viewing, (2) *Friends* viewing, (3) free recall, (4) post-movie picture viewing, and (5) viewing of shorter clip of a *Friends* episode for training a shared response model (SRM). Data from phases 1 and 4 were not reported in this paper. Additionally, T1 and T2 anatomical scans were conducted before and after all phases, respectively.

### MRI collection parameters and preprocessing

Neuroimaging data were acquired on a 3T full-body Siemens Skyra scanner with a 32-channel head coil, using a T2*-weighted echo planar imaging (EPI) pulse sequence (simultaneous multislice factor 3, TR 2000 ms, flip angle 67 degrees, TE 24 ms, whole-brain coverage 72 slices with a resolution of 1.9×1.9×2 mm, FOV 180 mm). The first preprocessing steps were performed using FMRIprep ^157^, including motion correction, susceptibility distortion correction, brain tissue segmentation, and coregistration and affine transformation of the functional volumes to the 1-mm isotropic T1w anatomical (TR = 1900 ms; TE = 3.1 ms; field of view = 256 mm^2^; flip angle = 7 degrees; image matrix = 256 x 256, 208 axial slices with 1.0 mm^3^ voxel) and subsequently to MNI space.

The data were imported and re-sampled to a 2-mm isotropic resolution aligned to the 2-mm MNI152 template. Next, the data were spatially smoothed using SUSAN smoothing with a 2-mm full width-half maximum spatial kernel. Next, the data were masked using an across-run averaged mask, z-scored, and confound variables [movement in three directions, rotation in three directions, framewise displacement, and six anatomical components used to correct for the influence of physiological noise ^158^] were regressed out at the run level. To account for hemodynamic lag for pattern similarity and neural event state analyses, we shifted time series data by 2 TRs (4 s).

### Regions of interest (ROIs) and cortical parcellation analyses

We were primarily interested in activity from six ROIs. V1 was created using voxels near the calcarine sulcus with the strongest inter-subject correlation in a prior study ^159^. mPFC was calculated using whole-brain functional connectivity, separating the default mode network into 10 parts with significant correlations and labeling the masks based on cluster location ^47^. AG was created by combining PGp and PGa ^39^ from the Julich brain atlas ^160^. PMC was taken from a prior study ^40^ using an atlas based on resting state functional connectivity ^161^. postHC was created by combining the HC body and tail and antHC from the HC head from a prior study ^162^.

To investigate across-cortical effects, we extracted activity from 100 parcels using the Schaefer atlas ^101^ and divided them into 7 subnetworks ^163^.

### Event pattern similarity analysis

To compute pattern similarity across events, we first averaged the BOLD activity within each voxel in an ROI across all time-shifted TRs of an event to create an event pattern. Then, we computed Pearson correlations between the patterns of each pair of events, after which we Fisher *z*-transformed the correlation values. This became the two-dimensional event similarity matrix shown in Figure 3B.

Next, we computed three multiple regression models, each run separately for each participant, with across-event similarity for each ROI as the outcome variable. The following variables were predictors in Models 1 and 2: time [in which we calculated the average correlation between an event and each other event at that specific event lag (e.g., +1, +2) to account for the autocorrelated BOLD signal]; storyline (1 for events within the same storyline, 0 otherwise); semantic (based on cosine similarity from the Universal Sentence Encoder ^99^) between the transcript of each pair of events); characters (based on the combination of characters in common across events); and places (based on whether events shared the same place or not). In Model 1, we added the mean causal matrix as another predictor, whereas in Model 2, we added the top causal matrix. In Model 3, which was conducted to predict the presence of storyline effects in the absence of predictors based on causality, we used the time and storyline predictors only.

### Event middle-boundary match analyses

To compute neural similarity between the middle and boundaries of events, we began by defining the middle as 4 TRs (8s) after event onset until 4 TRs before its boundary and the boundary as 2 TRs before the boundary until 2 TRs after (Figure 4A). Note we also conducted a separate analysis with only the exact boundary TR ^36^ (see Results). Defining middle events using these timings required dropping 10 (out of 52) shorter events. We then found the average event pattern for each voxel in each ROI, after which we correlated and Fisher *z*-transformed the pattern between each event middle and boundary (boundary-middle match). We removed events within ±2 events from the current event boundary from the analysis to avoid spurious values stemming from the autocorrelated BOLD signal (Figure 4B). Next, we computed inferential statistics on three different boundary-middle match contrasts for each participant within each ROI. First, we subtracted all past minus all future boundary-middle matches (Figure 4C). Next, we subtracted only past minus future boundary-middle matches with a non-zero value from the mean causal matrix (Figure 4D-E). Finally, we also subtracted causal minus non-causal past events (Figure 4F). Another analysis involved matching the middle event pattern from a given participant with the average event boundary pattern from all other participants, followed by similar inferential statistics (see Results). This analysis provided yet another control for concerns about the autocorrelated BOLD signal and, because it produced similar results, it suggests that event and event boundary templates are shared across individuals.

### Greedy state boundary search (GSBS) & shared response model (SRM) training

Following prior approaches ^39,86,164^, we used SRM ^105^ to reduce the dimensionality of the neural data to make neural event state analyses more computationally tractable. This involved finding patterns shared across participants given some number of features, *k*. The optimal *k* is unknown; we chose it by examining the best alignment with event boundaries in an independent video stimulus. First, we applied an across-participant SRM to each ROI from the viewing of an independent stimulus (an 11.5 min clip from another episode of *Friends*), iterating over a range of *k* values (min: 5, max: 40) by using GSBS to find neural event boundaries (max number of event states: 100, or approximately 8.7 / minute of stimulus) and examining its alignment with experimenter-defined boundaries from that stimulus. Next, we chose *k* based on which one had the highest correlation rank when measured against 10,000 iterations of the correlation against a time course of experimenter-defined events with scrambled event orders. For this analysis, we applied light smoothing to the neural (±2 TRs) and experimenter-defined event boundary time courses (±1 TR).

Next, we applied SRM to our main stimulus using the optimal *k* for each ROI. Then, we ran GSBS on SRM-based activation values for each ROI in each subject. On the main stimulus, we used a maximum of 350 event states (approximately 8.3 / minute of stimulus).

### Alignment between behavioral and neural event boundaries and effects of causality

First, we verified the alignment between three different behavioral event boundaries: experimenter-defined boundaries, agreement from a first group of participants performing event segmentation (EBA1), and agreement from a second group of participants performing event segmentation (EBA2) (Figure 5A).

Using outputs from GSBS applied to our main stimulus (Figure 5B), we aligned neural event boundaries with the three behavioral event boundaries (Figure 5C). For our neural event boundary time course, we ran GSBS on each participant independently and averaged the proportion of participants with event boundaries at each TR. We then smoothed both behavioral and neural time courses using a running mean of ±1 TR. Then, we correlated the time courses and Fisher *z*-transformed the values.

To examine the effects of causal network distance between successive events, we first quantified the mean neural event boundary strength surrounding each event boundary (±3 TRs surrounding each boundary) for each participant (Figure 5D). Next, we correlated this time course against the weighted path length between events in the causal network. We also repeated this identical analysis using neural event boundary likelihood as a binary (1 for boundary, 0 for no boundary) rather than boundary strength.

### Statistical analyses

To assess whether there was more evidence for the top causal strategy than other strategies, we computed within-participant *t*-tests between evidence for the top causal strategy against evidence for each other strategy independently (Figure 2D).

To assess whether causal or top causal models predicted event-by-event neural similarity based on multiple regression analyses, we examined whether the distribution of betas for each predictor differed from zero using one-sample *t*-tests separately for each ROI.

For the boundary–middle match analyses, we computed each subtracted boundary–middle match first for each participant and then compared this distribution to zero using one-sample *t*-tests separately for each ROI (Figure 4C,E-F). To address issues in the case of timing differences between the causal versus non-causal past events (wherein causally related events were closer to the current event on average), we used bootstrapping. Here, we randomly sampled boundary-middle matches with replacement 10,000 times and only examined contrasts when there were no significant differences (*p >*= 0.5) in the number of TRs between the randomly selected events. Next, we examined the proportion of the time the distribution of boundary-middle matches that were above zero, with significance indicated by the true value landing outside the central 95% (2.5-97.5th percentile).

To assess whether there was significant alignment between neural and non-neural event boundaries, we found the rank of the true Fisher *z*-transformed value against a distribution of values after running 10,000 iterations of the same correlation against a time course of experimenter-defined events with scrambled event orders (as indicated by the violin-outlined distributions in Figure 5C). We considered true values significant if they landed outside the middle 95% range (2.5-97.5th percentile).

To compare neural event boundary strength against causal network weighted path length, we Fisher *z*-transformed the correlation values across events and examined whether they differed from zero across participants for each ROI using one-sample *t*-tests (Figure 5E).

All analyses using cortical parcellations were conducted similarly to the above, except we plotted them according to significance values using uncorrected *p* values as well as false discovery rate (FDR)-corrected *q* values using the Benjamini/Hochberg method for independent or positively correlated tests ^165^.

The analyses in this study were not pre-registered.

## Code and data availability

All relevant code and data necessary to reproduce these results are available at https://github.com/JamesWardAntony/friends upon publication. Raw neuroimaging data are available on https://openneuro.org/datasets/ds008464 in the brain imaging data structure (BIDS) format.

## Acknowledgements

The following individuals assisted with data collection and scoring: Himanshu Chaudhary, Mitchell A. Nguyen, Ashley Monteiro, and Hannah Pries. We also acknowledge funding from the Office of Naval Research (ONR) Multidisciplinary University Research Initiatives (MURI) Program (N00014-17-1-2961) and the National Science Foundation (NSF) Behavioral and Cognitive Sciences Division (BCS) (2122550).

## Author Contributions

J.W.A. conceived, programmed, and analyzed all data from the experiments, and drafted the manuscript. S.A. contributed to data collection from the event segmentation experiments. M.S.B. helped with transcription and recall scoring from the fMRI recall data. Z.M.R. contributed analysis ideas. C.R. helped conceive and analyze the experiments and contributed heavily to editing the paper.

## Competing interests statement

The authors declare no competing interests.

## Supplementary tables and figures

**Figure S1.**
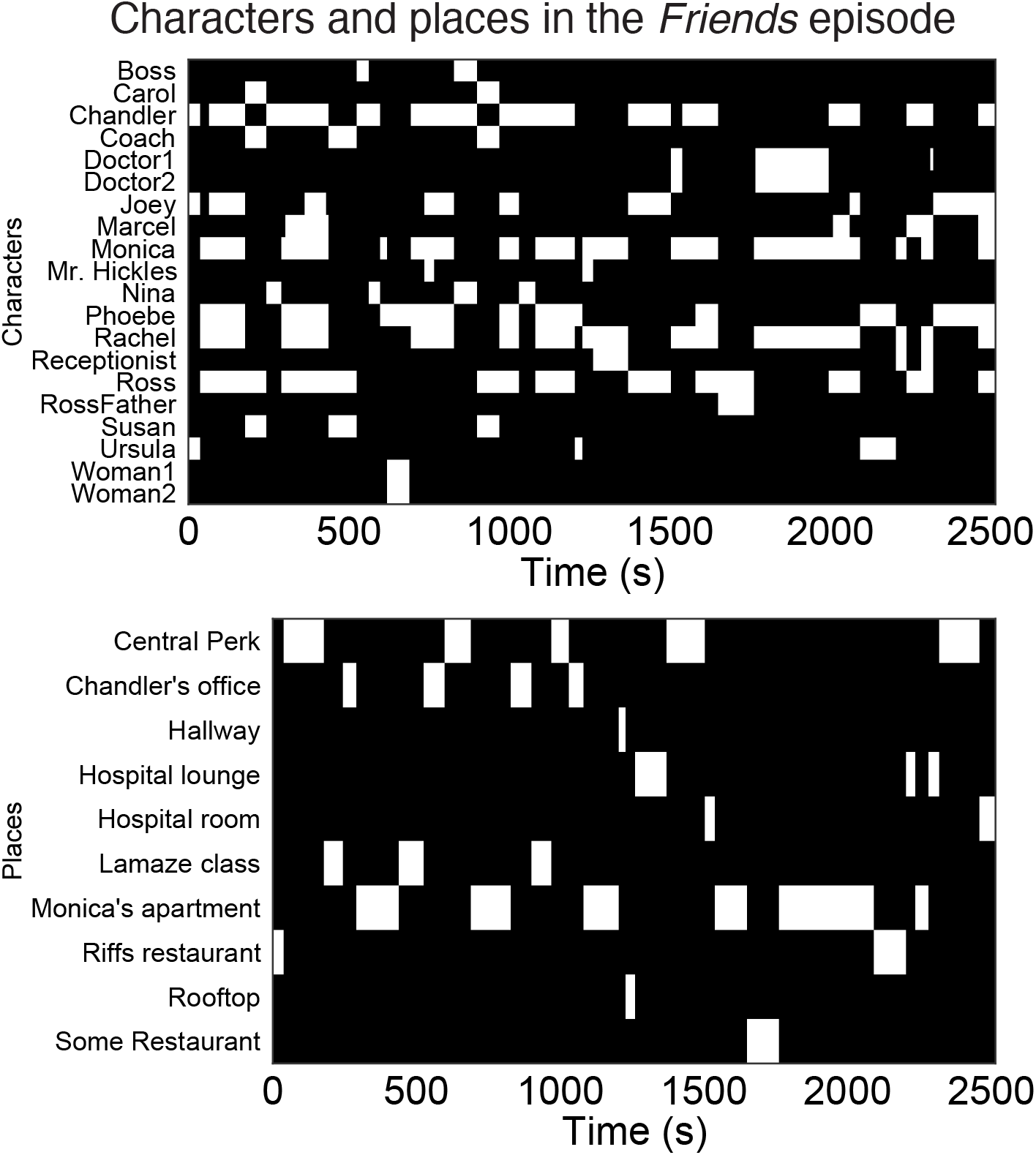
Timeline of characters and places in the two-part *Friends* episode. Related to Figure 1.

**Figure S2.**
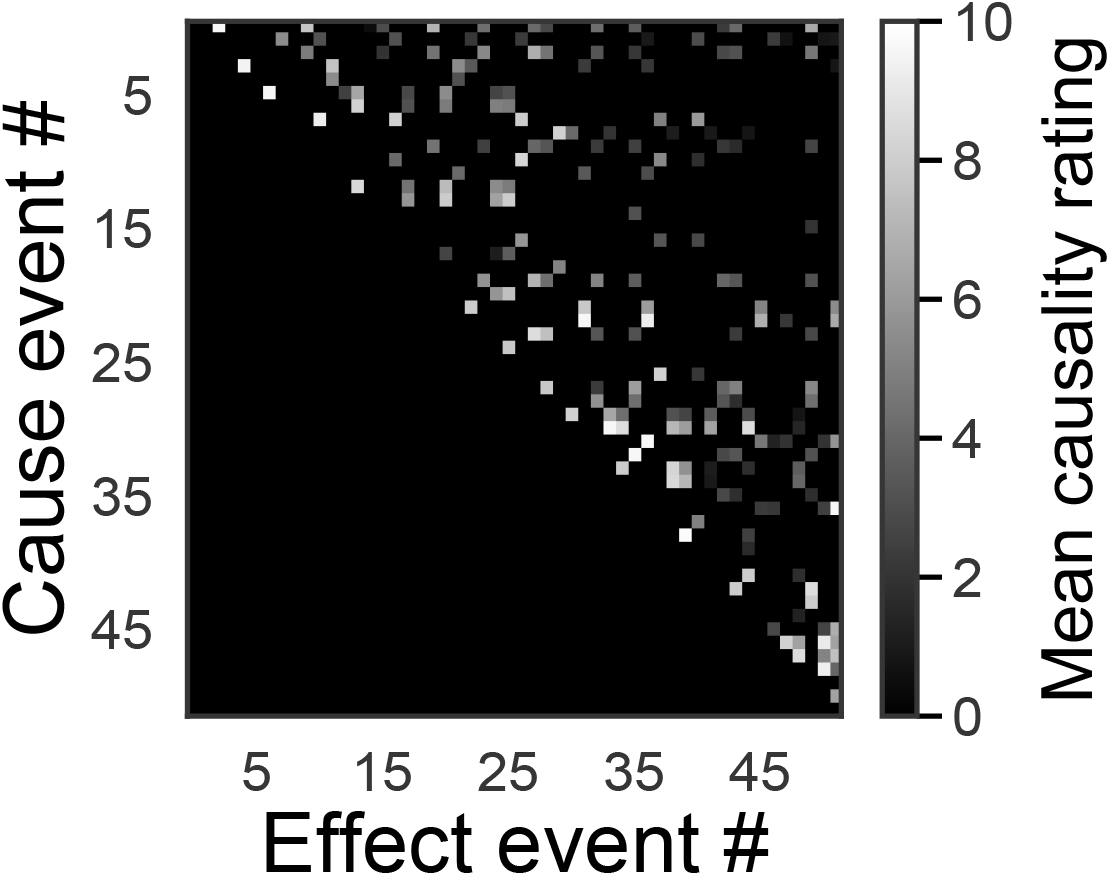
Mean across-rater causality values, from which the network in Figure 1C was determined. Related to Figure 1.

**Figure S3.**
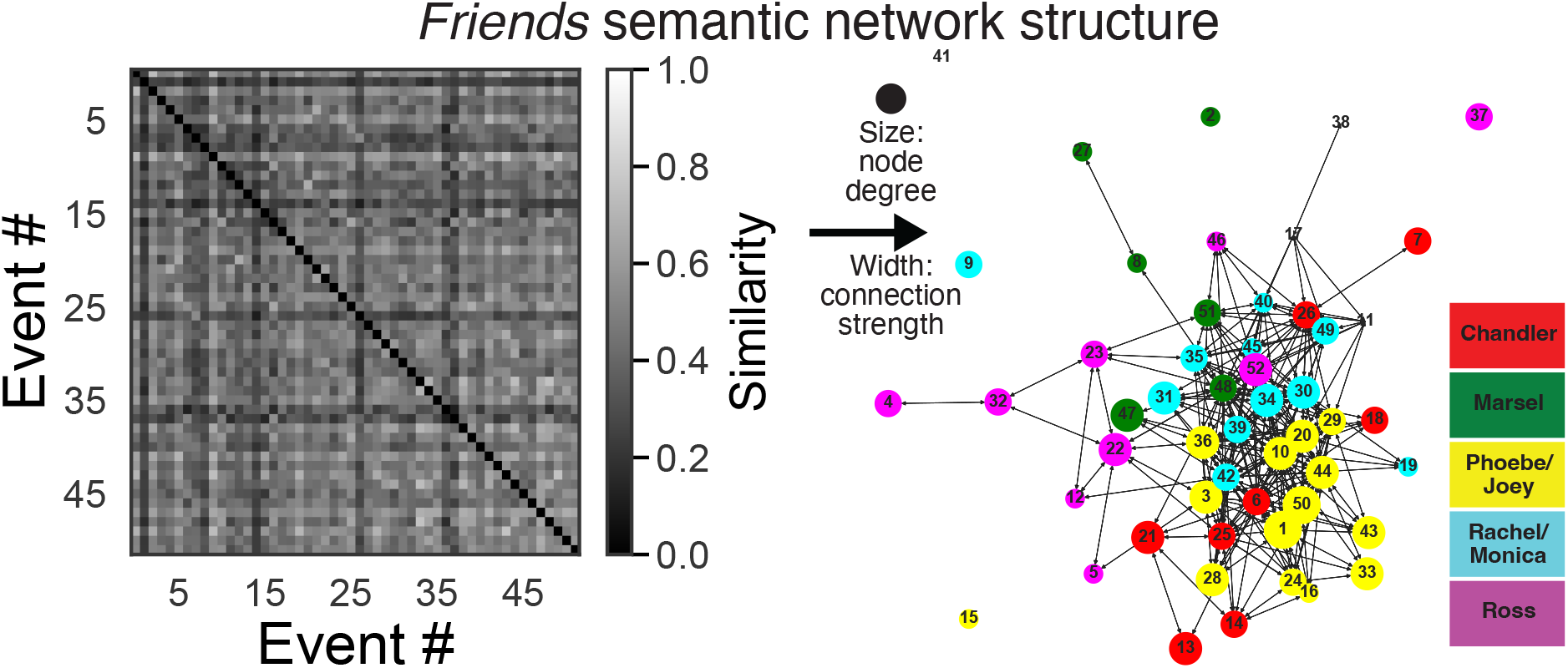
(left) Semantic similarity between events based on the cosine similarity between the text of the show transcript using the Universal Sentence Encoder ^99^. (right) Semantic structure based on these similarity values. Related to Figure 1.

**Figure S4.**
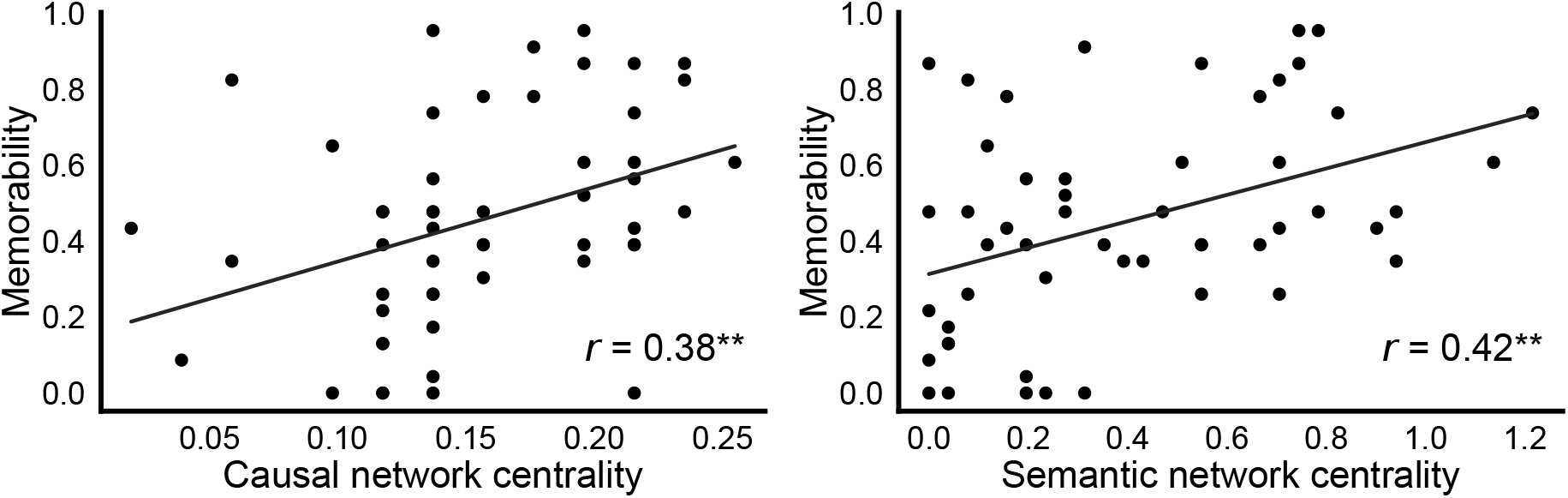
Event memorability (proportion of participants recalling each event) correlates with centrality (node degree) in both causal (left) and semantic networks (right). These measurements predict *what* participants remember, whereas the focus of the main manuscript is on *how* they remember, or how they transition between events. Related to Figure 2.

**Figure S5.**
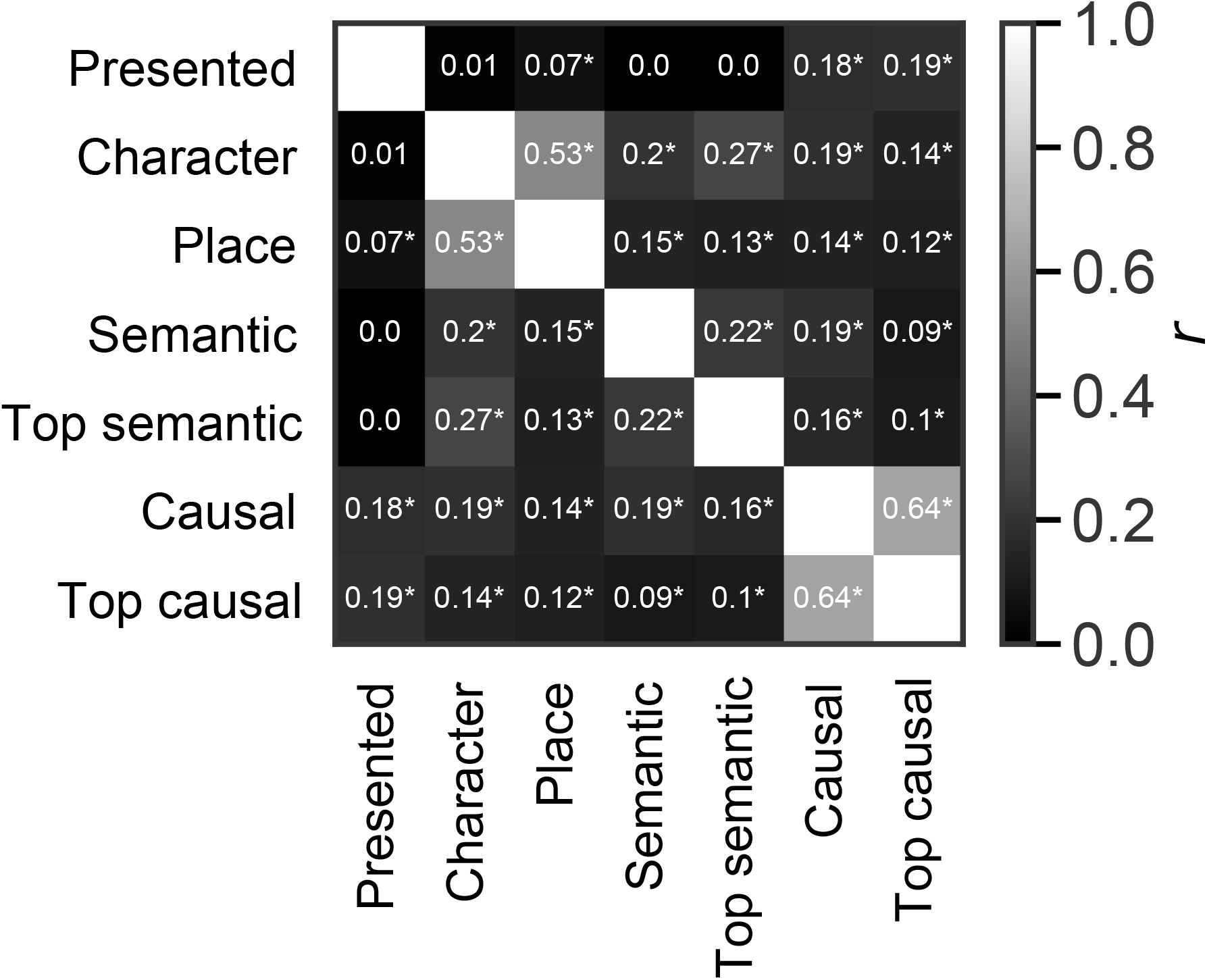
Correlation coefficients among all strategy transition matrices. Asterisks indicate *p* < 0.05. Related to Figure 2.

**Figure S6.**
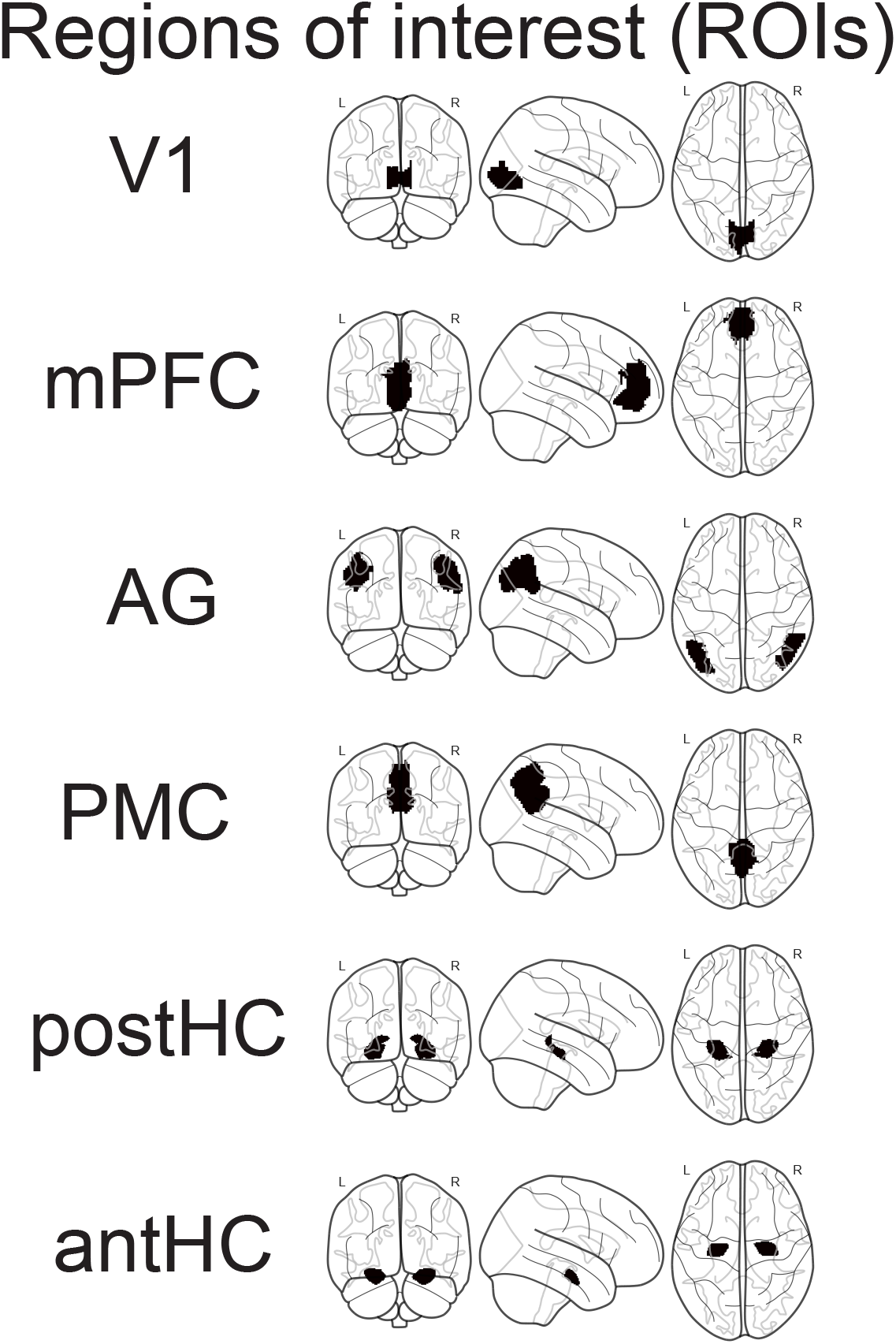
Six ROIs plotted on glass brain. Related to Figure 3, 4, 5.

**Figure S7.**
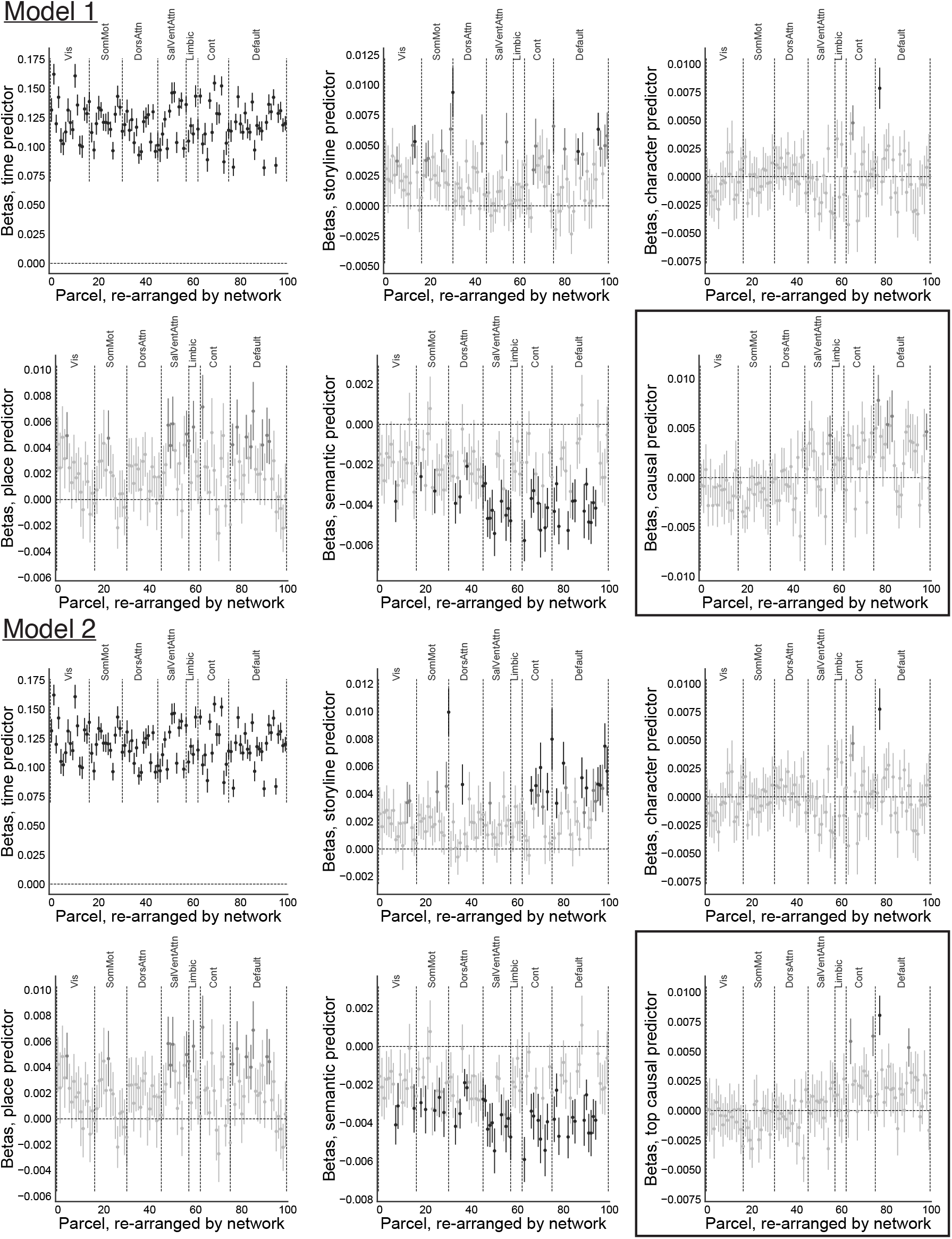
Across-parcel results from Models 1-2 grouped by seven broader networks ^163^, with parcels in black indicating significance after FDR-correction for all parcels (*q <* 0.05), in dark gray indicating uncorrected significance (*p* < 0.05), and all non-significant parcels in light gray. Related to Figure 3.

**Figure S8.**
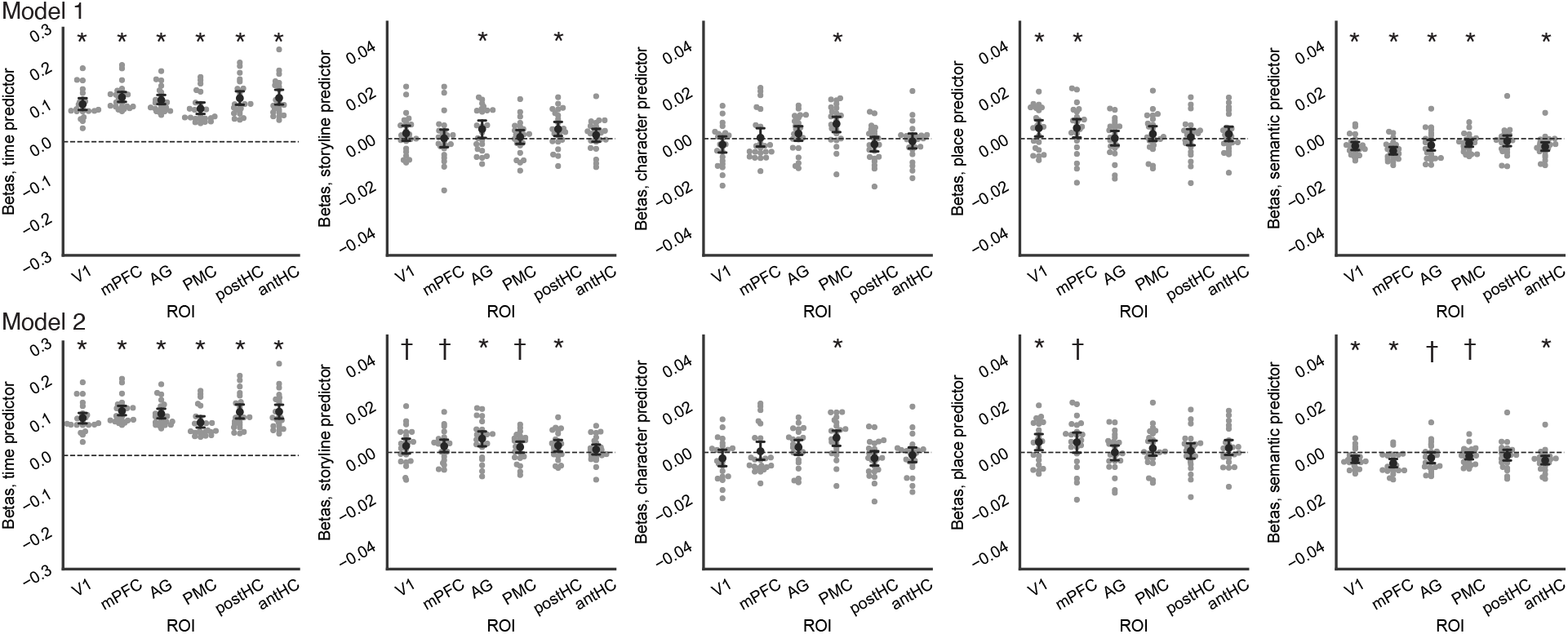
Results from Model 1 and Model 2 for all non-causal predictors. Related to Figure 3.

**Figure S9.**
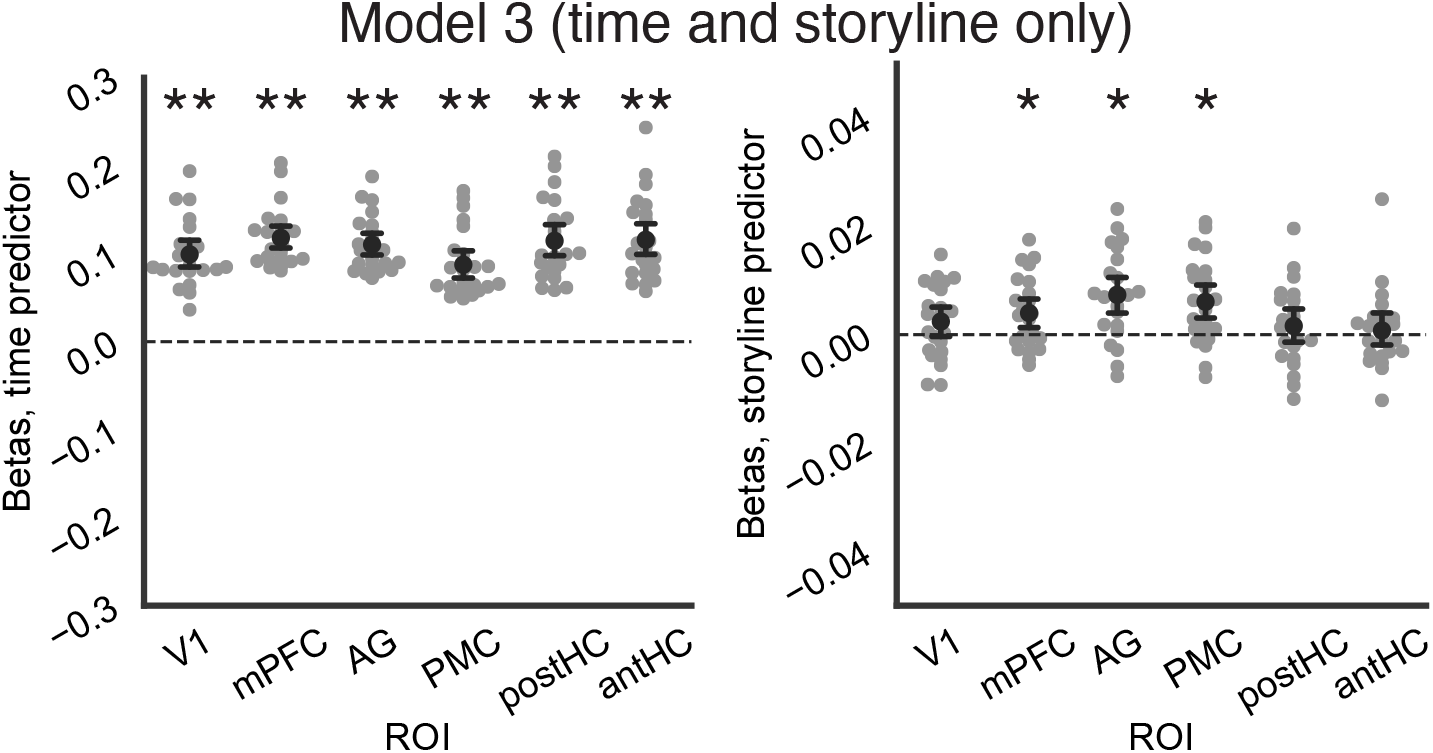
Results from Model 3 with only time and storyline as predictors. Related to Figure 3.

**Figure S10.**
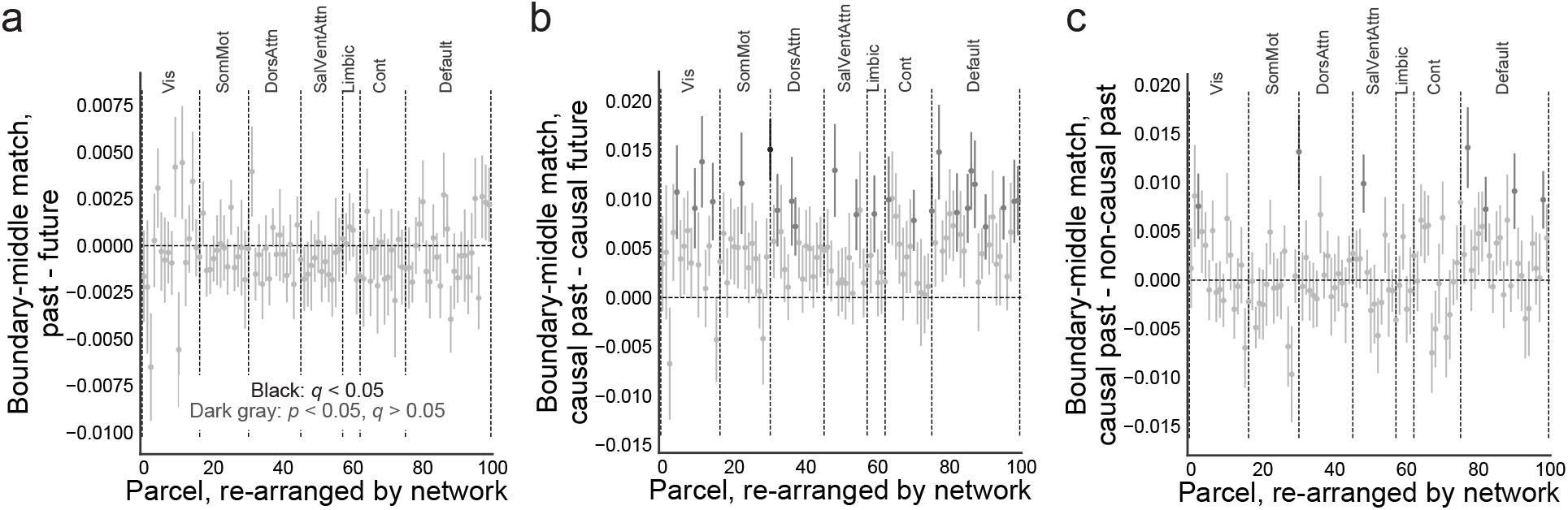
Across-parcel boundary-middle match analyses for all past minus future (A), causally related past minus future (B), and causal minus non-causal past (C). Results were grouped by seven broader networks ^163^, with parcels in black indicating significance after FDR-correction for all parcels (*q <* 0.05), in dark gray indicating uncorrected significance (*p* < 0.05), and all non-significant parcels in light gray. Related to Figure 4.

**Figure S11.**
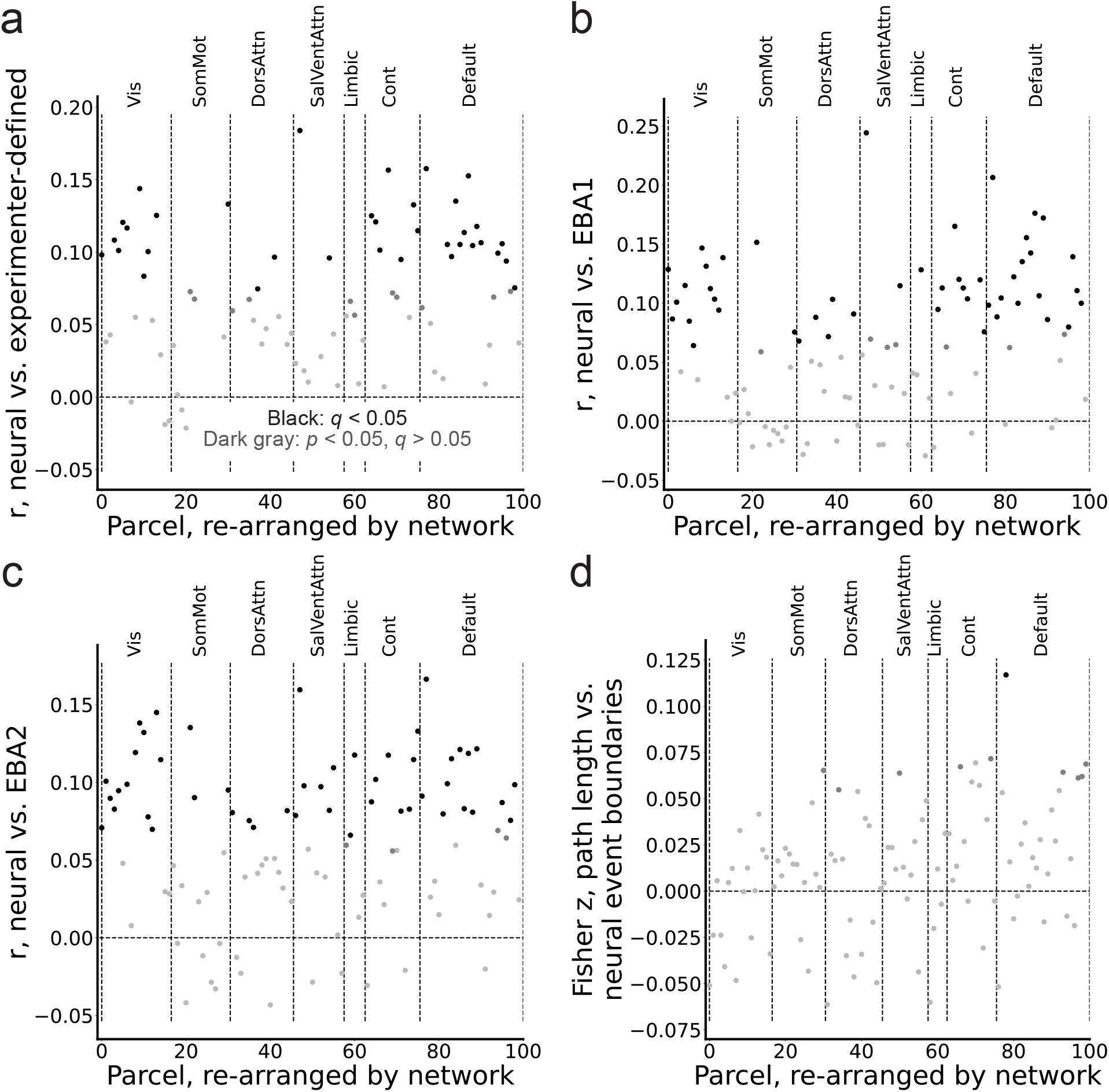
Across-parcel results showing how neural event boundaries aligned with experimenter-defined boundaries (A), event boundary agreement from the first and second segmentation experiments (EBA1/EBA2) (B-C), and how well event boundary strength correlated with weighted path lengths within the causal network (D). Results were grouped by seven broader networks ^163^. *P* values were initially determined by rank against identical correlations between neural and behavioral event boundaries after scrambling event orders. After this, black depicts significant parcels after FDR-correction (*q <* 0.05), dark gray depicts uncorrected significance (*p* < 0.05), and light gray depicts all non-significant parcels. Related to Figure 5.

## Notes

### Competing Interest Statement

The authors have declared no competing interest.

### Summary of Updates

Basic changes to title and abstract.

